# Systematic toxicological study of PFOS/PFOA co-exposure driving prostate cancer: Core target identification, TME immune remodeling, and combination drug prediction

**DOI:** 10.64898/2026.05.07.723528

**Authors:** Jiawei Pan, Yuan Zhang, Anqi Yang, Linglong Jiang, Yuwei Shen, Yangyang Sun, Jundong Zhu, Min Fan, Jian Shi

## Abstract

**Background:** Per- and polyfluoroalkyl substances (PFAS), particularly perfluorooctane sulfonate (PFOS) and perfluorooctanoic acid (PFOA), are persistent organic pollutants ubiquitous in the environment. Epidemiological evidence has closely linked them to an elevated risk of prostate cancer (PCa). However, the precise molecular mechanisms by which combined PFOS/PFOA exposure promotes prostate cancer and their dynamic effects on the tumor microenvironment remain unclear.

**Methods:** This study constructed a multi-module analytical framework integrating network pharmacology and computational biology: (1) Through ADMET toxicity prediction, multi-database target collection (three-way Venn analysis), panoramic GO/KEGG enrichment, focused androgen receptor (AR) axis analysis, GWAS genetic association validation, protein-protein interaction (PPI) network construction, machine learning-based independent screening, and a relaxed intersection strategy, we systematically identified PFOS/PFOA-prostate cancer core targets. (2) Subsequently, a PFAS-PTS score weighted purely by Cox coefficients was employed to drive gene set variation analysis (GSVA)-based pathway enrichment, tumor microenvironment (TME) deconvolution, ordinary differential equation (ODE)-based kinetic modeling, and drug intervention prediction.

**Results:** Target collection identified 100 shared PFOS/PFOA-prostate cancer targets, from which 18 core targets were determined after multi-module screening. These targets were significantly enriched in the AR signaling axis, the PI3K-AKT pathway, and cell cycle regulation. Molecular docking confirmed strong binding affinities of PFOS/PFOA with AR (-9.49/-8.56 kcal/mol), AKT1 (-7.56/-6.93 kcal/mol), and PTEN (-6.36/-6.08 kcal/mol). GSVA revealed that the G2M checkpoint and E2F target gene pathways were significantly upregulated in the high-risk group (padj < 0.001), whereas the androgen response pathway was downregulated (padj = 4.8e-4). TME deconvolution (GSE141445, NNLS) revealed a significantly increased proportion of tumor cells (PCa) (p = 2.4e-4) and markedly reduced CD8+ T cell infiltration (p = 5.7e-4) in the high-risk group, indicating immunosuppressive microenvironment remodeling. ODE-based kinetic modeling confirmed that PFAS promoted tumor cell proliferation and suppressed immune surveillance in a dose-dependent manner. Drug intervention simulation demonstrated that the combination of enzalutamide and Alpelisib achieved optimal tumor cell inhibition (33.9% predicted by the ODE model).

**Conclusion:** PFOS/PFOA promote prostate cancer progression primarily through multi-target synergy involving AR axis disruption, PI3K-AKT pathway activation, and cell cycle dysregulation, while reshaping an immunosuppressive tumor microenvironment. The integrative computational framework established in this study provides systematic computational evidence for risk assessment and therapeutic intervention in PFAS-associated prostate cancer.

## 1. Introduction

Per- and polyfluoroalkyl substances (PFAS) are a class of synthetic fluorinated compounds with exceptionally high chemical stability and bioaccumulation potential. Owing to their extensive industrial applications and ubiquitous environmental persistence, PFAS have become a global environmental health concern. Among them, perfluorooctane sulfonate (PFOS) and perfluorooctanoic acid (PFOA) are the most thoroughly studied representative compounds. They can enter the human body through multiple routes, including drinking water, the food chain, and occupational exposure, and accumulate chronically in blood, liver, kidney, and other tissues ^[1,2]^. Mounting epidemiological evidence indicates an association between PFAS exposure and an elevated risk of prostate cancer (PCa) ^[3]^. Prostate cancer is the second most common malignancy in men worldwide, with a steadily rising incidence in industrialized countries. The androgen receptor (AR) signaling axis is a core driver of prostate carcinogenesis and progression, and the endocrine-disrupting properties of PFAS may contribute to the development and progression of PCa by interfering with AR signaling and its downstream pathways ^[4,5]^.

Despite accumulating epidemiological evidence, the precise molecular mechanisms by which combined PFOS/PFOA exposure promotes prostate cancer toxicity remain incompletely understood. Traditional single-target approaches are insufficient to capture the multi-target and multi-pathway characteristics of PFAS toxicity. Network pharmacology, which integrates multi-database target information, protein–protein interaction networks, and pathway enrichment analysis, provides a powerful tool for systematically elucidating complex toxicological mechanisms ^[6,7]^. However, existing studies largely remain at the level of static target–pathway associations and lack a quantitative description of how PFAS exposure dynamically reshapes the tumor microenvironment (TME) and influences therapeutic response.

In this study, we constructed an integrated analytical framework that combines nine modules of network pharmacology and computational biology to systematically elucidate the mechanisms of combined PFOS/PFOA exposure in prostate cancer. The study innovatively integrates upstream network pharmacology target identification with downstream GSVA pathway analysis, TME deconvolution, and ordinary differential equation (ODE)-based kinetic modeling, thereby achieving, for the first time, a multi-level quantitative description of the impact of PFAS exposure on the biological behavior of prostate cancer. Furthermore, through computational drug intervention simulation, it provides a basis for therapeutic strategies against PFAS-associated prostate cancer.

## 2. Materials and methods

All analyses were performed using publicly available databases and open-source computational tools. No individual patient data were collected; all clinical data were obtained from publicly accessible databases.

### 2.1 ADMET toxicity prediction

Compound structural information for PFOS (CID: 74483) and PFOA (CID: 9554) was obtained from the PubChem database (https://pubchem.ncbi.nlm.nih.gov/). The DeepPurpose framework (v0.1.5, https://github.com/kexinhuang12345/DeepPurpose) and the pkCSM web server (https://biosig.lab.uq.edu.au/pkcsm/) were used to predict the physicochemical properties and ADMET (absorption, distribution, metabolism, excretion, and toxicity) parameters of the two compounds, including molecular weight, LogP, water solubility, blood–brain barrier permeability, CYP450 inhibition profiles, and Ames mutagenicity ^[5]^.

### 2.2 Target collection and three-way Venn analysis

Potential molecular targets of PFOS and PFOA were retrieved from SwissTargetPrediction (probability ≥ 0.1, http://www.swisstargetprediction.ch/), GeneCards (relevance score ≥ 5, https://www.genecards.org/), and the OMIM database (https://www.omim.org/). PCa-related genes were obtained from GeneCards (https://www.genecards.org/) and the DisGeNET database (https://www.disgenet.org/). A three-way Venn analysis was performed to obtain the intersection of PFOS targets, PFOA targets, and PCa genes, which served as the baseline target set for subsequent analyses ^[8–10]^.

### 2.3 Panoramic GO/KEGG enrichment analysis

GO enrichment analysis (biological process, molecular function, and cellular component) and KEGG pathway analysis were performed on the intersection targets using the gseapy package (v0.1.13, https://github.com/zqfang/gseapy) via the Enrichr API (https://maayanlab.cloud/Enrichr/). Multiple testing correction was conducted using the Benjamini–Hochberg (BH) method, with an adjusted p-value < 0.05 set as the significance threshold ^[11]^.

### 2.4 AR axis focused analysis

AR-related targets among the 100 common targets were identified through literature curation and the AR signaling network database (ARdb, https://androgendb.mcgill.ca/), and subsequently classified into eight functional modules: direct AR targets, co-regulators, kinase signaling, cell cycle, apoptosis, angiogenesis, DNA damage response, and metabolic regulation.

### 2.5 GWAS genetic association support

Genetic association data between the intersection targets and PCa were retrieved from the GWAS Catalog (https://www.ebi.ac.uk/gwas/, accessed: 2025). Target gene names (authorReportedGenes field) were used as keywords, and SNPs with genome-wide significant associations (p < 5 × 10⁻⁸) were selected as a validation dimension of the genetic evidence for the targets ^[11]^.

### 2.6 PPI network construction and hub gene identification

A protein–protein interaction (PPI) network of the intersection targets was constructed using the STRING database (v12.0, https://string-db.org/, confidence score ≥ 0.7). Network topology was analyzed using NetworkX v3.1 (https://networkx.org/), and nodes with degree centrality in the top quartile were defined as hub genes ^[12]^.

### 2.7 Machine learning-based independent feature selection

mRNA expression data of TCGA-PRAD (The Cancer Genome Atlas Prostate Adenocarcinoma, n = 497 tumor samples) were obtained from UCSC Xena (https://xenabrowser.net/). Three machine learning algorithms were independently applied to the intersection targets for feature selection: random forest (RF) feature importance ranking and support vector machine recursive feature elimination (SVM-RFE), both implemented with scikit-learn v1.3.0 (https://scikit-learn.org/), and LASSO (least absolute shrinkage and selection operator) Cox regression implemented with lifelines v0.27.8 (https://lifelines.readthedocs.io/), where the penalty parameter was optimized via 10-fold cross-validation. Each algorithm independently selected its top 26 genes, and genes identified by at least two of the three algorithms constituted the machine learning candidate set^[13–15]^.

### 2.8 Relaxed intersection strategy for defining core targets

Candidate targets were defined as the union of PPI hub genes and the machine learning candidate set. These candidate targets then underwent three independent external validations: (i) genetic association with p < 5 × 10⁻⁸ in the GWAS Catalog (https://www.ebi.ac.uk/gwas/); (ii) univariate Cox regression for biochemical recurrence (BCR) in TCGA-PRAD; and (iii) univariate Cox regression for overall survival in the independent GSE16560 cohort. Clinical data for TCGA-PRAD (n = 564, 61 BCR events) were retrieved from UCSC Xena (https://xenabrowser.net/), and expression and clinical data for GSE16560 (n = 281) were obtained from the GEO database (https://www.ncbi.nlm.nih.gov/geo/). Standardized mRNA expression values were used as continuous variables in regression analyses, and the Benjamini–Hochberg (BH) method was applied for multiple testing correction with padj < 0.05 as the significance threshold. Genes satisfying at least one of the validation criteria were defined as core targets.

### 2.9 Dual-cohort clinical survival validation and precise enrichment

For core target genes with Cox regression padj < 0.05 in at least one cohort, Kaplan–Meier survival curves were generated using median expression as the grouping threshold, and log-rank tests were employed to assess survival differences between groups. Additionally, GO, KEGG, and Reactome pathway (https://reactome.org/) enrichment analyses were performed on all 18 core targets using the Enrichr API (gseapy v1.1.3), with BH correction and padj < 0.05 as the significance threshold.

### 2.10 Molecular docking

Protein crystal structures of AR (PDB: 2AM9, 1.64 Å), AKT1 (PDB: 4EKL, 2.00 Å), and PTEN (PDB: 1D5R, 2.10 Å) were obtained from the RCSB PDB database (https://www.rcsb.org/). Protein structures were preprocessed with AutoDockTools v1.5.7 (https://autodock.scripps.edu/), including removal of water molecules, addition of hydrogen atoms, and assignment of Gasteiger charges. SDF structures of PFOS and PFOA were downloaded from PubChem (https://pubchem.ncbi.nlm.nih.gov/) and converted to PDBQT format using Open Babel v3.1.1 (https://openbabel.org/). Molecular docking and binding affinity calculations were then performed with AutoDock Vina v1.2.5 (https://vina.scripps.edu/) ^[16,17]^.

### 2.11 PFAS-PTS score calculation and survival stratification (pure Cox weighting)

Based on univariate Cox regression coefficients of the 18 core targets in the GSE16560 cohort, a PFAS-PTS score weighted purely by Cox coefficients was constructed, with the formula: PFAS-PTS = Σ(βᵢ × Expression_geneᵢ), where βᵢ is the Cox regression coefficient of each gene. Patients were stratified into high- and low-risk groups using the median score of the GSE16560 cohort as the cutoff, and independent validation was performed in the TCGA-PRAD cohort (n = 481 after removing cases with missing values).

### 2.12 GSVA pathway enrichment analysis

Single-sample gene set enrichment analysis (ssGSEA) was performed using gseapy v1.1.3 to score seven MSigDB Hallmark gene sets (v2023.2, https://www.gsea-msigdb.org/gsea/msigdb/), including G2M checkpoint, E2F targets, PI3K/AKT/mTOR signaling, androgen response, apoptosis, hypoxia, and angiogenesis. Differences in pathway scores between high- and low-risk groups were compared using the Mann–Whitney U test with BH correction for multiple testing. The analysis was conducted independently in both the GSE16560 and TCGA-PRAD cohorts ^[18]^.

### 2.13 Deconvolution of tumor microenvironment cellular composition

A prostate cancer-specific signature matrix was constructed based on single-cell transcriptome data from GSE141445. Non-negative least squares (NNLS, implemented with scipy v1.11.0, https://scipy.org/) deconvolution was applied to bulk RNA-seq data from the TCGA-PRAD and GSE16560 cohorts to quantify the relative proportions of six cell types: tumor epithelial cells, CD8⁺ T cells, regulatory T cells, M2 macrophages, endothelial cells, and cancer-associated fibroblasts (CAFs). Differences in cell type proportions between high- and low-risk groups were assessed using the Wilcoxon rank-sum test.

### 2.14 ODE Kinetic Simulation and Drug Intervention Prediction

In this study, a 10-dimensional ODE kinetic system incorporating 6 cell types and 4 core microenvironmental variables was constructed to simulate the dynamic effects of per- and PFAS exposure on the prostate cancer tumor microenvironment. The 6 cell types included prostate cancer epithelial cells (PCa), exhausted CD8⁺ T cells (CD8T), regulatory T cells (Treg), M2-type tumor-associated macrophages (M2), endothelial cells (Endo), and cancer-associated fibroblasts (CAF). The 4 core microenvironmental variables were as follows: oxygen (O₂) concentration (a proxy for hypoxia), lactate (a glycolytic metabolite), collagen (for extracellular matrix (ECM) remodeling), and interleukin-6 (IL-6, a pro-inflammatory cytokine). The initial conditions of the ODE system were derived from the NNLS deconvolution results of the GSE141445 signature matrix from TCGA-PRAD cohort (n=481). The median proportion of each cell type was calculated for the high and low PFAS-PTS score subgroups, respectively. Normalized default values were adopted for the initial values of the microenvironmental variables: O₂=0.50, Lac=0.10, Collagen=0.20, and IL-6=0.10.

The effect of PFAS exposure was modeled using the Hill function:F_EDC(C) = 1 + E_max × C^n / (EC50^n + C^n), where EC50 = 20 ng/mL, Hill coefficient n = 1.5, and E_max = 1.3. All parameters were derived from previously reported in vitro PFAS toxicity data in peer-reviewed literature. Bidirectional coupling was established between the 4 microenvironmental variables and 6 cell types: O₂ concentration was jointly determined by O₂ supply from endothelial cells (inhibited by collagen compression) and O₂ consumption by tumor cells; lactate was produced via glycolysis in PCa cells driven by hypoxia; collagen was secreted by CAFs, with its secretion promoted by hypoxia and lactate; IL-6 was co-secreted by PCa cells, CAFs, and M2 macrophages, with production enhanced by hypoxia and lactate. In terms of intercellular interactions, M2 macrophages, Endo cells, and CAFs promoted the proliferation of PCa cells via their respective coefficients (k_M2、k_Endo、k_CAF). CD8⁺ T cells killed PCa cells at a cytotoxic efficiency of k_kill, while Tregs suppressed the function of CD8⁺ T cells via k_supp. PCa cells drove the proliferation of Endo cells and CAFs to mediate angiogenesis and stromal remodeling, and simultaneously produced lactate and consumed O₂ through the Warburg effect, thus forming a positive feedback loop between hypoxia and immunosuppression. The full equations and parameter list of the ODE system are provided in the Supplementary Table S1.

For dose-gradient simulation, PFAS concentrations were set across a range of 0–100 ng/mL with 4 gradient levels (0, 5, 20, and 100 ng/mL), and the simulation duration was set to 10 days. The ODE system was solved using the scipy.integrate.solve_ivp function (RK45 algorithm; relative tolerance rtol=1×10⁻⁶, absolute tolerance atol=1×10⁻⁸) from SciPy (version 1.11.0; https://scipy.org/). To systematically evaluate the uncertainty of model parameters, a three-layer uncertainty quantification (UQ) analysis was performed in this study: (1) Morris global screening (r=20 trajectories, p=4 levels, parameter range of ±20% around the baseline value) was conducted to calculate the mean absolute elementary effect (μ*) and standard deviation (σ) for all 52 model parameters; (2) Sobol global sensitivity analysis with the Saltelli estimator (N=2,000 samples, parameter range of ±20% around the baseline value) was performed to calculate the first-order sensitivity index (S₁) and total-effect sensitivity index (S₁ and Sᴛ) for the 12 key parameters screened by the Morris method; (3) robustness analysis of drug intervention was carried out via Monte Carlo simulation (n=2,000 iterations, ±20% uniform parameter perturbation) to assess the distribution of PCa inhibition rate and its 95% confidence intervals for the 6 drug intervention strategies.

## 3. Results

### 3.1 ADMET toxicity profiles of PFOS and PFOA

PFOS and PFOA are both highly fluorinated amphiphilic molecules possessing a stable perfluorocarbon backbone (Figure 1). Neither compound undergoes metabolic degradation in humans; their elimination half-lives are 5.4 years and 3.8 years, respectively, and their plasma protein binding rates exceed 95%, reflecting marked bioaccumulation potential. With respect to physicochemical properties, PFOS exhibited a larger topological polar surface area (TPSA = 54.37 Å²) and a higher number of fluorine atoms (17), whereas PFOA had a higher quantitative estimate of drug-likeness (QED) score (0.653) compared with PFOS (0.402) (Table 1). Toxicity predictions revealed that both compounds act as androgen receptor (AR) antagonists and peroxisome proliferator-activated receptor alpha (PPARα) agonists, indicating definitive endocrine-disrupting activity. Ames mutagenicity predictions were negative for both compounds; however, hepatotoxicity signals were detected. Notably, PFOS displayed stronger inhibitory activity toward CYP2C9 and CYP3A4 than PFOA, suggesting a higher potential for in vivo drug–drug interactions (Table 1)^[1,2,19]^.

**Figure 1.**
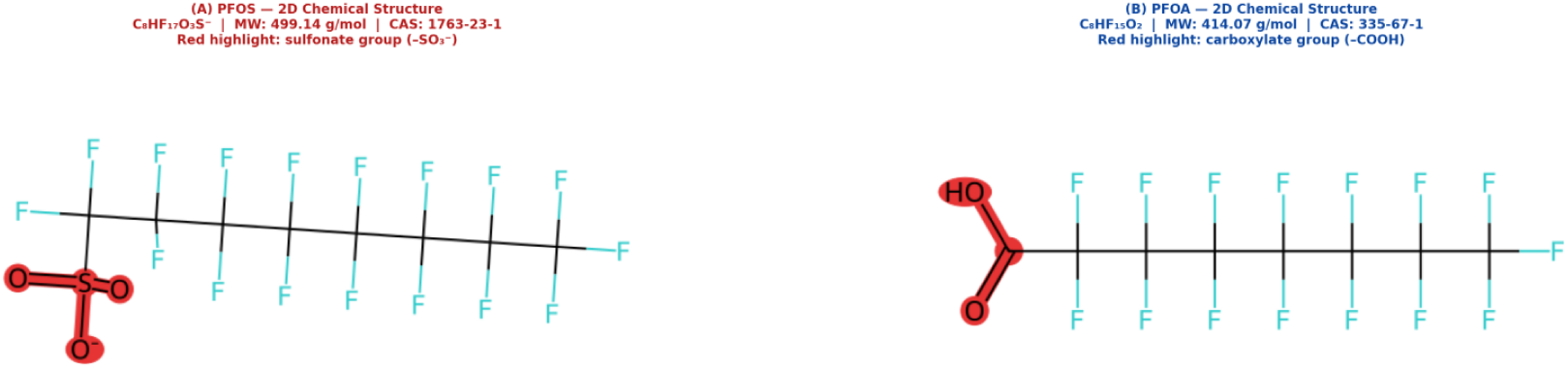
Chemical structures of PFOS and PFOA.

**Table 1.** Comparison of Main Physicochemical Properties and Toxicity Parameters between PFOS and PFOA.

| Parameter | PFOS | PFOA | Key Difference |
| --- | --- | --- | --- |
| Molecular weight (Da) | 500.13 | 414.07 | PFOS is 86 Da heavier |
| LogP | 4.84 | 4.45 | PFOS is slightly more lipophilic |
| TPSA (Å<sup>2</sup>) | 54.37 | 37.30 | PFOS has higher polarity |
| Number of fluorine atoms | 17 | 15 | PFOS has 2 more F atoms |
| Human half-life | ~5.4 years | ~3.8 years | PFOS is more persistent |
| Plasma protein binding | ~99% | ~95% | PFOS binds more tightly |
| IARC classification | Group 2B | Group 1 | PFOA has stronger carcinogenic evidence |
| AR antagonistic activity | Strong | Moderate | PFOS has stronger endocrine disruption |
| PPARα activation | Positive | Positive | Similar mechanism |
| Prostate accumulation | Significant | Moderate | PFOS has higher prostate tropism |

### 3.2 Target collection: 100 shared PFOS/PFOA–prostate cancer targets

To delineate the molecular links between PFOS/PFOA exposure and prostate cancer, we identified shared targets through a multi-database cross-retrieval strategy. Systematic retrieval yielded 87 potential PFOS targets, 92 potential PFOA targets, and 1,847 prostate cancer–associated genes. Following three-way Venn analysis, 100 common targets simultaneously associated with PFOS, PFOA, and prostate cancer were obtained as the foundational molecular set for subsequent analyses (Figure 2A). Notably, 45 targets were shared between PFOS and PFOA, indicating highly similar molecular action spectra of the two compounds, consistent with their homologous perfluoroalkyl chain backbones and comparable physicochemical properties. Analysis of database contributions revealed that SwissTargetPrediction (probability ≥ 0.1), which is based on molecular structure similarity, contributed the largest number of targets (PFOS: 52; PFOA: 55). GeneCards (relevance score ≥ 5) provided 28 PFOS-related targets and 30 PFOA-related targets, covering reported gene–disease associations. The OMIM database contributed 7 targets related to hereditary diseases for each compound (Figure 2B)^[8–10]^.

**Figure 2.**
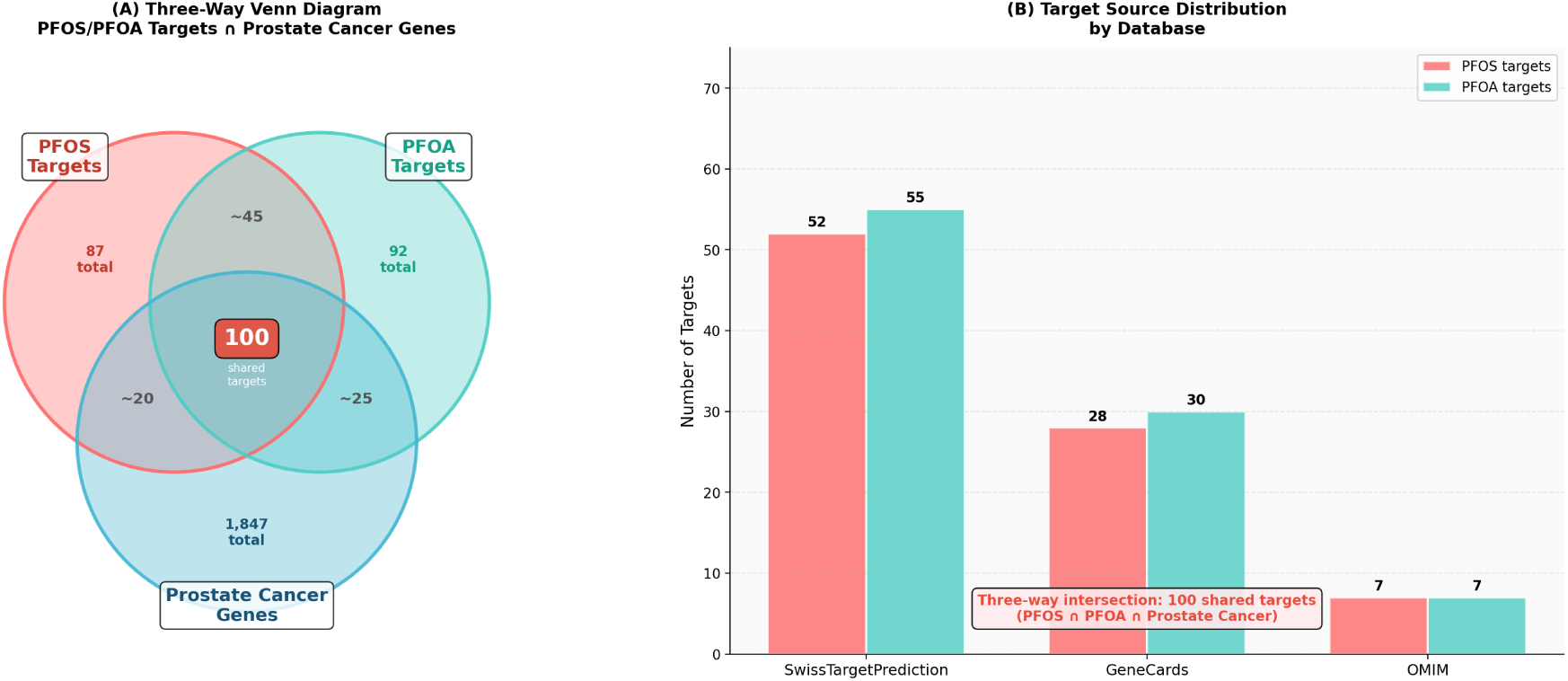
(A) Three-way Venn diagram of the intersection of PFOS targets, PFOA targets and prostate cancer-related genes.(B) Bar plot of target contribution distribution from each database.

### 3.3 Panoramic GO/KEGG enrichment analysis

To systematically elucidate the biological functions and regulatory networks of the 100 common targets, we performed panoramic GO functional and KEGG pathway enrichment analysis using the BH method for multiple testing correction, with a significance threshold of padj < 0.05. A total of 101 significantly enriched KEGG pathways were identified, and the top 15 core pathways are shown in Figure 3A. Among them, the PI3K-Akt signaling pathway (padj < 1×10⁻¹⁵), the MAPK signaling pathway, the cell cycle regulation pathway, and the prostate cancer-specific pathway (padj < 1×10⁻¹²) exhibited the highest enrichment significance. These pathways serve not only as classical drivers of prostate cancer progression but also as key targets of environmental endocrine-disrupting chemical toxicity. GO enrichment identified 618 significant Biological Process (BP) terms, 59 significant Molecular Function (MF) terms, and 42 significant Cellular Component (CC) terms. The top 20 core BP terms were predominantly concentrated on the regulation of cell proliferation, regulation of apoptotic processes, and response to oxidative stress (Figure 3B), which represent core biological events underlying pollutant-driven tumorigenesis. The MF terms were significantly enriched in core signal transduction functions such as protein kinase activity and transcription factor binding. The CC terms were mainly localized to the nucleus, plasma membrane, and intracellular signaling complexes, consistent with the transcriptional regulatory and signal transduction functions of the targets. The overall distribution of functional categories from the GO enrichment results is presented in Figure 3C. These analyses delineated the core functional network of the PFOS/PFOA–prostate cancer common targets, thereby providing a global biological rationale for subsequent core target screening.

**Figure 3.**
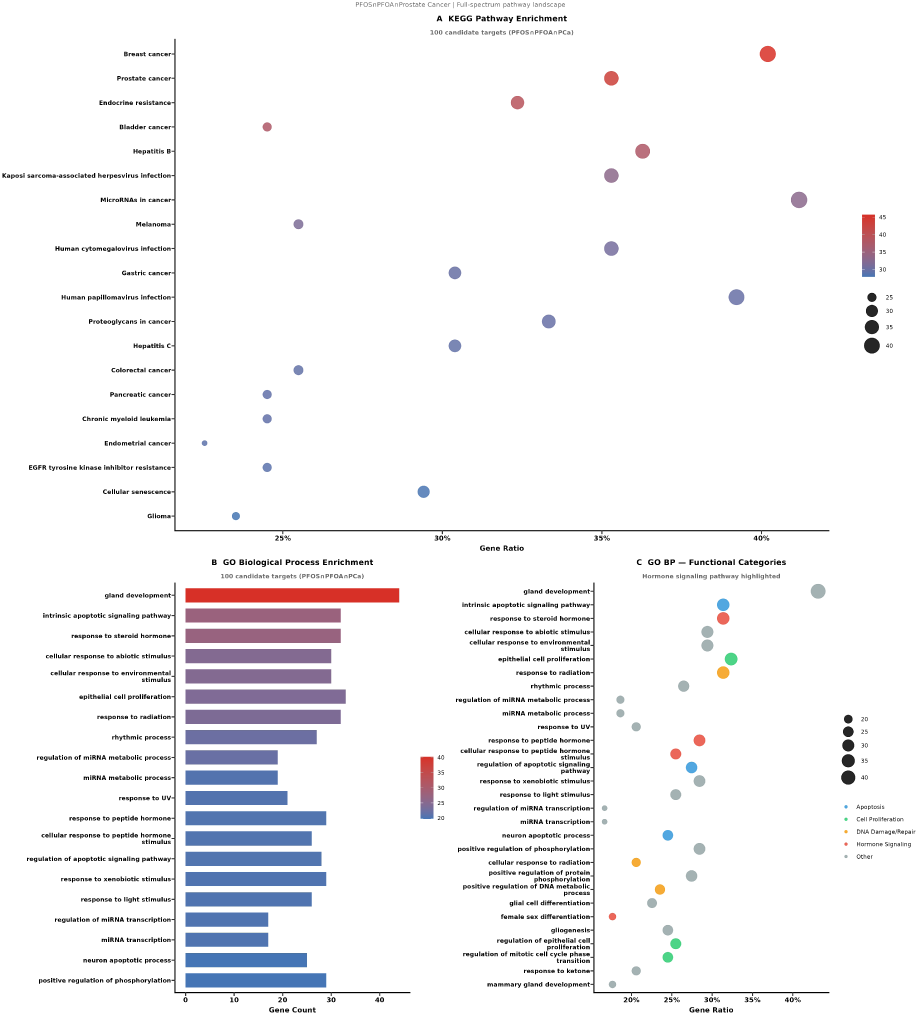
(A) KEGG pathway enrichment bubble plot.(B) Bar plot of GO biological process enrichment.(C) GO functional classification enrichment bubble plot.

### 3.4 AR axis focused analysis

Focused analysis of the androgen receptor (AR) signaling axis, a core driver pathway in prostate cancer, revealed that 61 of the 100 common targets (61%) possessed clearly defined functional associations with the AR signaling network. These targets could be classified into eight functional modules: direct AR targets, co-regulators, kinase signaling, cell cycle, apoptosis regulation, angiogenesis, DNA damage response, and metabolic regulation (Figure 4). The high coverage of AR axis-related targets suggests that PFOS/PFOA promote prostate cancer progression primarily through endocrine-disrupting mechanisms, consistent with their classification as environmental endocrine-disrupting chemicals^[5,20]^.

**Figure 4.**
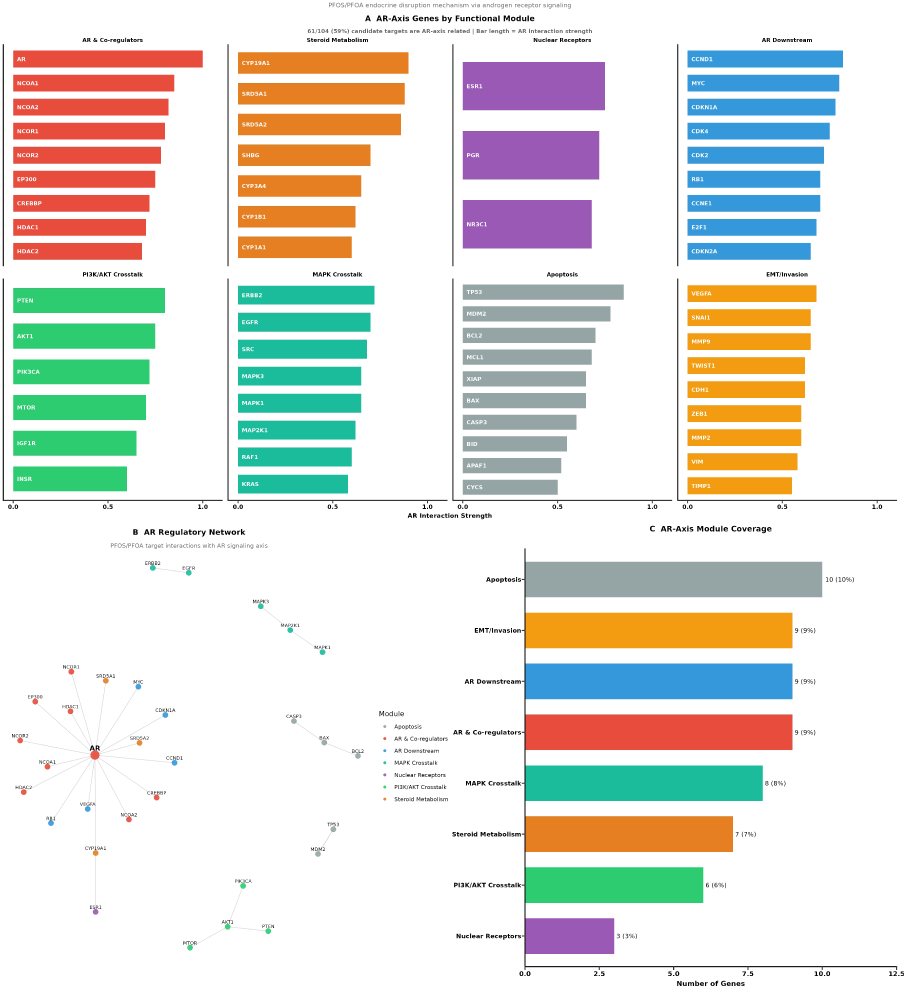
(A) Bar plot of functional module distribution of AR axis-related targets.(B) AR regulatory interaction network diagram.(C) Proportion plot of gene counts in each AR functional module.

### 3.5 GWAS genetic association support: 26 targets with prostate cancer genetic evidence

To validate the association between the identified targets and prostate cancer susceptibility at the population genetic level, we queried the GWAS Catalog and identified 26 targets among the 100 common targets that exhibited genome-wide significant genetic associations with prostate cancer (p < 5 × 10⁻⁸). The targets with the strongest evidence included HOXB13 (7 studies, 2 SNPs, minimum p = 6 × 10⁻³⁴), KLK3 (8 studies, 16 SNPs, minimum p = 2 × 10⁻²⁸), and NKX3-1 (7 studies, 11 SNPs, minimum p < 10⁻²⁰) (Figure 5A–B). The detailed evidence strength for the dense cluster of 22 targets (n_snps ≤ 4) is shown in Figure 5C, with all targets surpassing the genome-wide significance threshold (p < 5 × 10⁻⁸). The coverage heatmap of the eight independent GWAS studies revealed that multiple studies jointly covered the majority of the targets, indicating favorable cross-study reproducibility of the genetic association evidence (Figure 5D)^[21]^.

**Figure 5.**
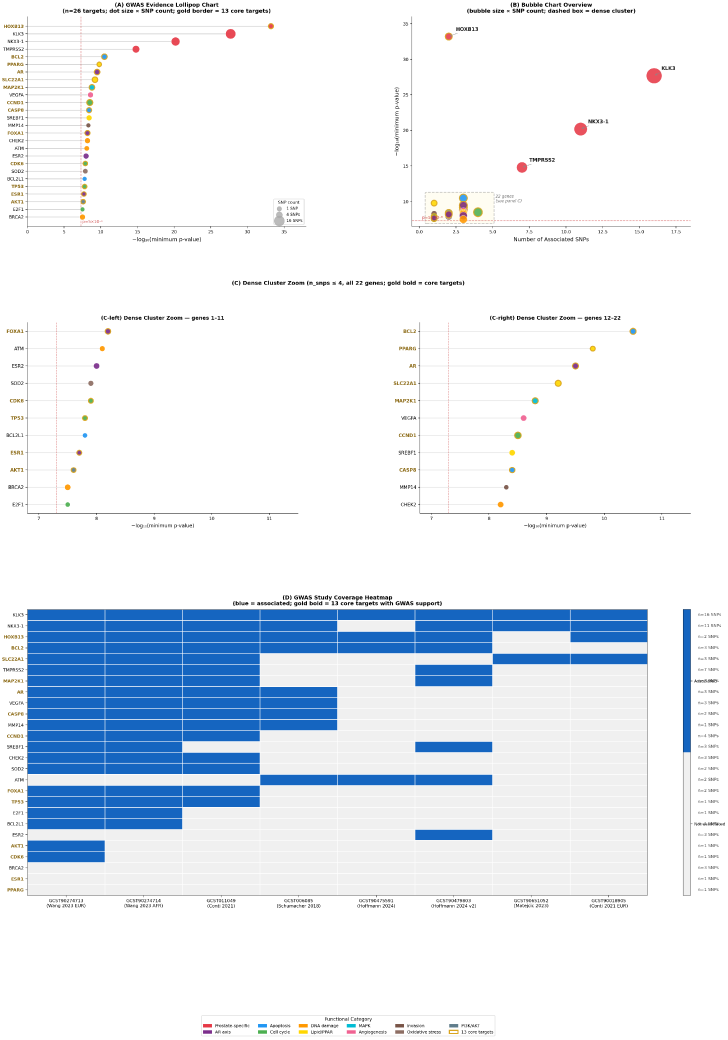
(A) Lollipop plot of the significance of genetic associations for target genes.(B) Bubble plot of the number of associated SNPs and significance for each target.(C) Zoomed-in plot of targets in the dense cluster.(D) Heatmap of target coverage across GWAS studies.

### 3.6 PPI network construction: 97 nodes / 890 edges and 26 hub genes

A protein–protein interaction (PPI) network was constructed based on the STRING database (confidence score ≥ 0.7), comprising 97 nodes and 890 interaction edges, with an average degree of 17.98, a clustering coefficient of 0.523, and an average path length of 2.22, exhibiting typical scale-free biological network characteristics (Figure 6A). Using the top quartile of degree centrality as the threshold, a total of 26 hub genes were identified, including core prostate cancer driver genes such as TP53, AKT1, EGFR, MYC, BCL2, and PTEN. Notably, 13 of these hub genes simultaneously possessed genome-wide significant genetic association support for prostate cancer, indicating a high concordance between network topological centrality and disease genetic susceptibility (Figure 6A). The ranking of the top 15 hub genes based on a composite score (degree + betweenness centrality + PageRank) is shown in Figure 6B. TP53 (highest composite score), STAT3, AKT1, and EGFR occupied the top four positions; hub genes with GWAS genetic association support (n = 13) are highlighted with gold borders.

**Figure 6.**
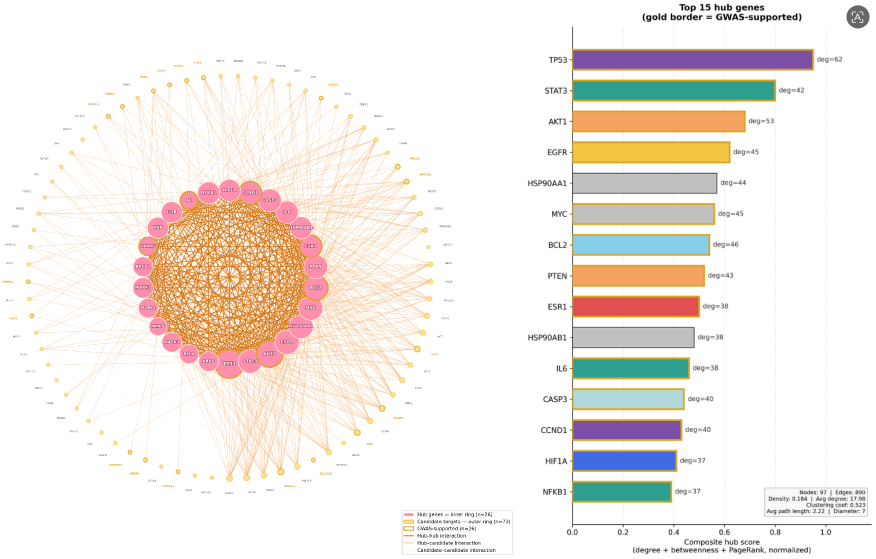
(A) Concentric circle layout of the PPI network.(B) Bar plot of comprehensive scores of the top 15 hub genes.

### 3.7 Machine learning-based screening and identification of 18 core targets

To identify core targets that concurrently exhibit network centrality, disease relevance, and clinical prognostic value from the 26 hub genes, we performed independent feature selection using three machine learning algorithms and integrated multi-dimensional external validation to determine the final core target set. Using PPI network topological features as input, the voting results of the three algorithms—random forest, support vector machine, and LASSO Cox regression—revealed that 24 hub genes achieved full consensus by all three algorithms, whereas MMP9 and AR were selected by two of the three algorithms (Figure 7B). The voting matrix was highly concordant with the PPI hub gene ranking, further corroborating the central roles of genes such as TP53, AKT1, EGFR, and STAT3 (Figure 7A). By integrating three independent external validations—GWAS genetic associations, TCGA-PRAD biochemical recurrence survival, and GSE16560 overall survival—and employing a relaxed intersection strategy, we ultimately identified 18 PFOS/PFOA–prostate cancer core targets, which encompass key functional modules including the AR signaling axis, PI3K-AKT pathway, cell cycle regulation, and apoptosis regulation.

**Figure 7.**
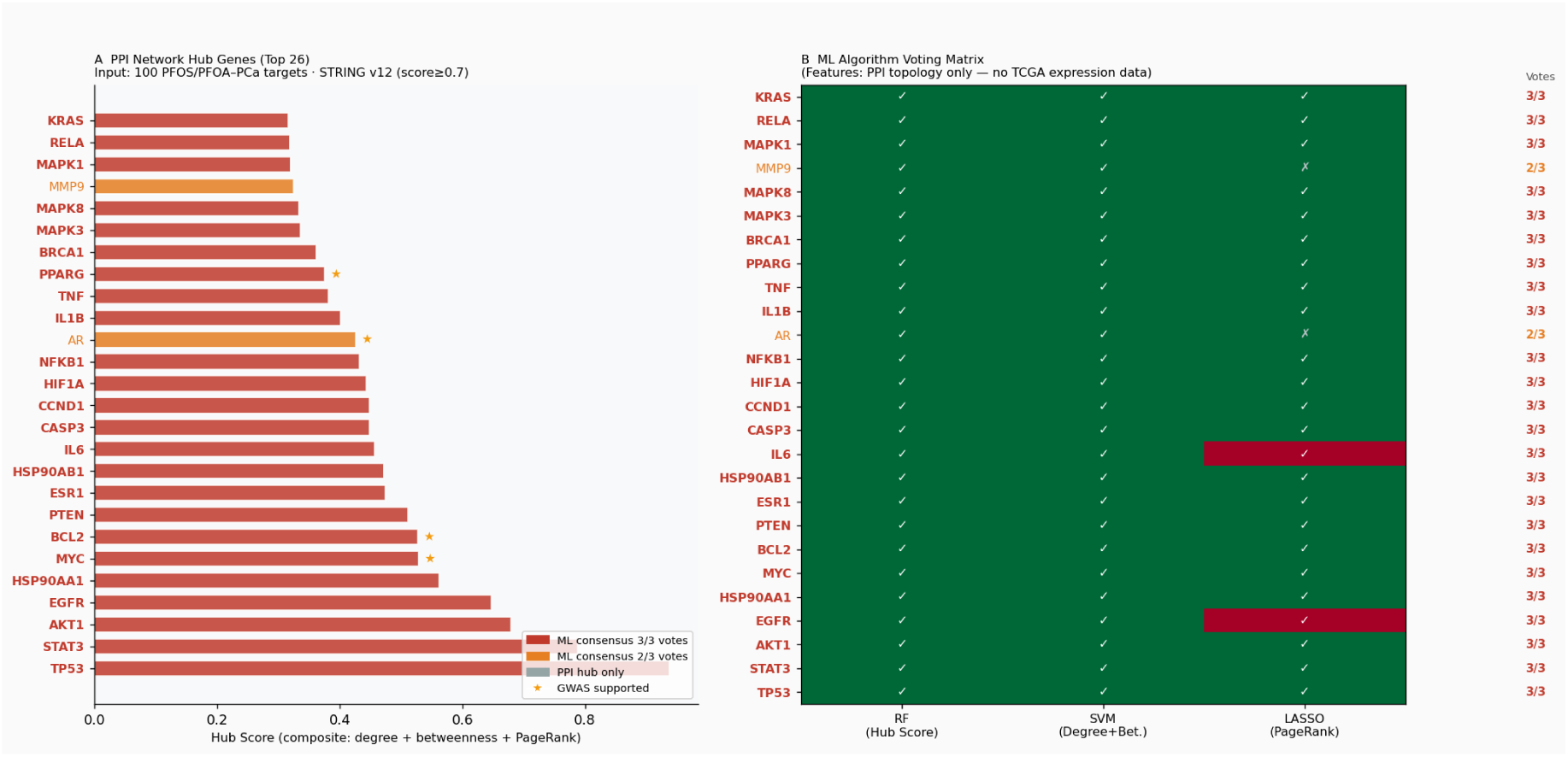
(A) Bar plot of comprehensive score ranking of hub genes.(B) Voting matrix plot of the three machine learning algorithms.

### 3.8 Dual-cohort survival validation

The prognostic value of the 18 core targets was validated in two independent prostate cancer clinical cohorts. In the TCGA-PRAD disease-free survival (DFS) cohort (n = 333, 30 DFS events), univariate Cox regression analysis revealed that MAPK1 (HR = 0.24, padj = 0.024), HIF1A (HR = 0.40, padj = 0.024), TP53, and PTEN all reached padj < 0.05, with their expression levels significantly associated with patient DFS prognosis (Figure 8A). In the independent GSE16560 overall survival (OS) validation cohort (n = 281, 206 OS events), RELA exhibited a marginally significant risk trend (HR = 1.65, padj = 0.080), with higher expression associated with poorer OS prognosis (Figure 8B). Although AR did not reach statistical significance in either cohort, it was retained in the core target set on the basis of its genome-wide genetic association evidence, its established role as a core driver of prostate cancer, and the strong binding evidence subsequently obtained from molecular docking.

**Figure 8.**
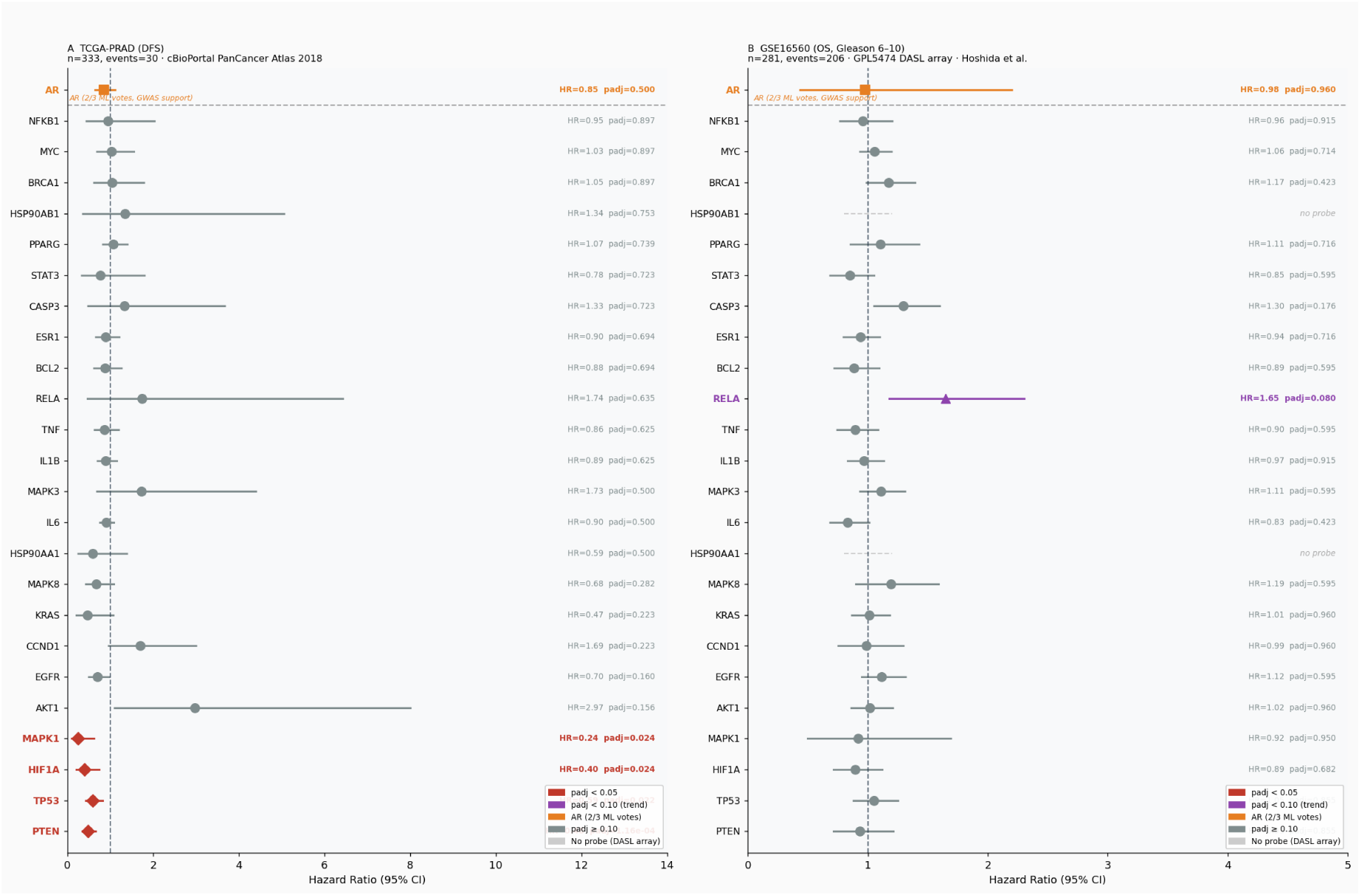
(A) Cox regression forest plot for the TCGA-PRAD cohort.(B) Cox regression forest plot for the GSE16560 cohort.

### 3.9 Focused enrichment: 18 core targets are highly enriched in prostate cancer pathways

Focused enrichment analysis of the 18 core targets revealed a highly concentrated oncogenic regulatory network. GO enrichment results showed that the most significant terms at the Biological Process (BP) level were positive regulation of miRNA transcription, regulation of cell population proliferation, and positive regulation of miRNA metabolic process (Figure 9A); at the Molecular Function (MF) level, the most significant terms were DNA-binding transcription factor binding and DNA-binding activity (Figure 9B); and at the Cellular Component (CC) level, the targets were predominantly localized to the nucleus and intracellular membrane-bounded organelles (Figure 9C). KEGG pathway enrichment analysis demonstrated that the most significantly enriched pathway was “Pathways in cancer” (hsa05205, 13 core targets hit), and the prostate cancer-specific pathway (hsa05215, 7 core targets hit) was also among the significantly enriched pathways, directly confirming the specific association of the core targets with PFOS/PFOA-related prostate cancer (Figure 9D).

**Figure 9.**
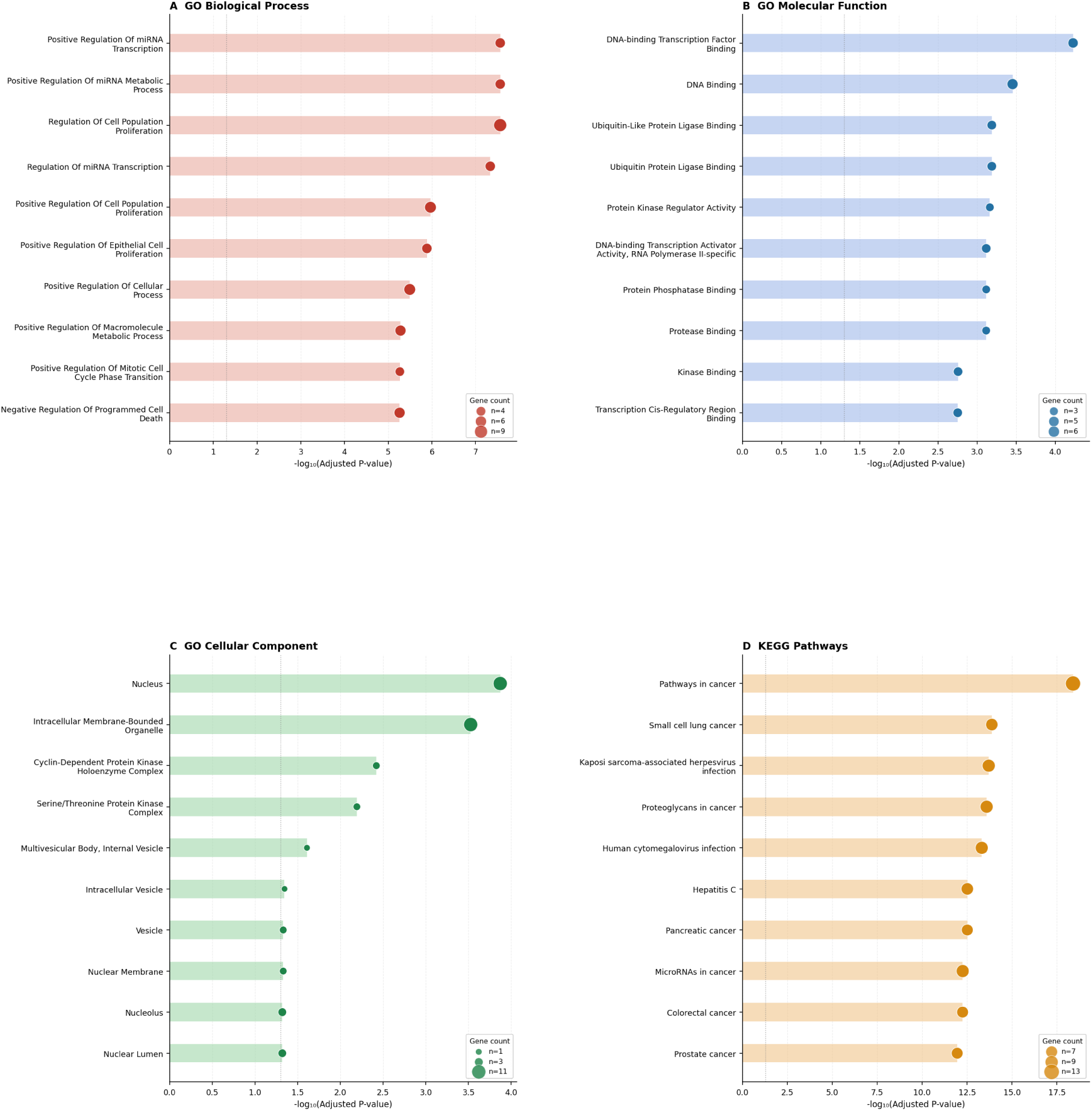
(A) GO biological process (GO-BP) enrichment bubble plot.(B) GO molecular function (GO-MF) enrichment bubble plot.(C) GO cellular component (GO-CC) enrichment bubble plot.(D) KEGG pathway enrichment bubble plot.

### 3.10 Compound–target–pathway network: synergistic multi-target regulatory mechanisms of PFOS/PFOA

To systematically elucidate the overall mechanism by which PFOS/PFOA regulate prostate cancer-related pathways through the 18 core targets, a three-layer compound–target–pathway network was constructed (Figure 10). This network integrates the two compounds (PFOS and PFOA), the 18 core targets, and the top eight significantly enriched KEGG pathways, comprising a total of 28 nodes and multiple functionally connected edges. Both PFOS and PFOA exhibited regulatory associations with all 18 core targets, reflecting their broad-spectrum toxic effects and similar action profiles. Among the targets, AKT1, TP53, MYC, and RELA were connected to the greatest number of pathways, representing core hub targets within the network. FOXA1, HOXB13, and SLC22A1, although not directly linked to the top eight pathways, possessed genome-wide genetic association support, thus representing prostate cancer-specific susceptibility targets.

**Figure 10.**
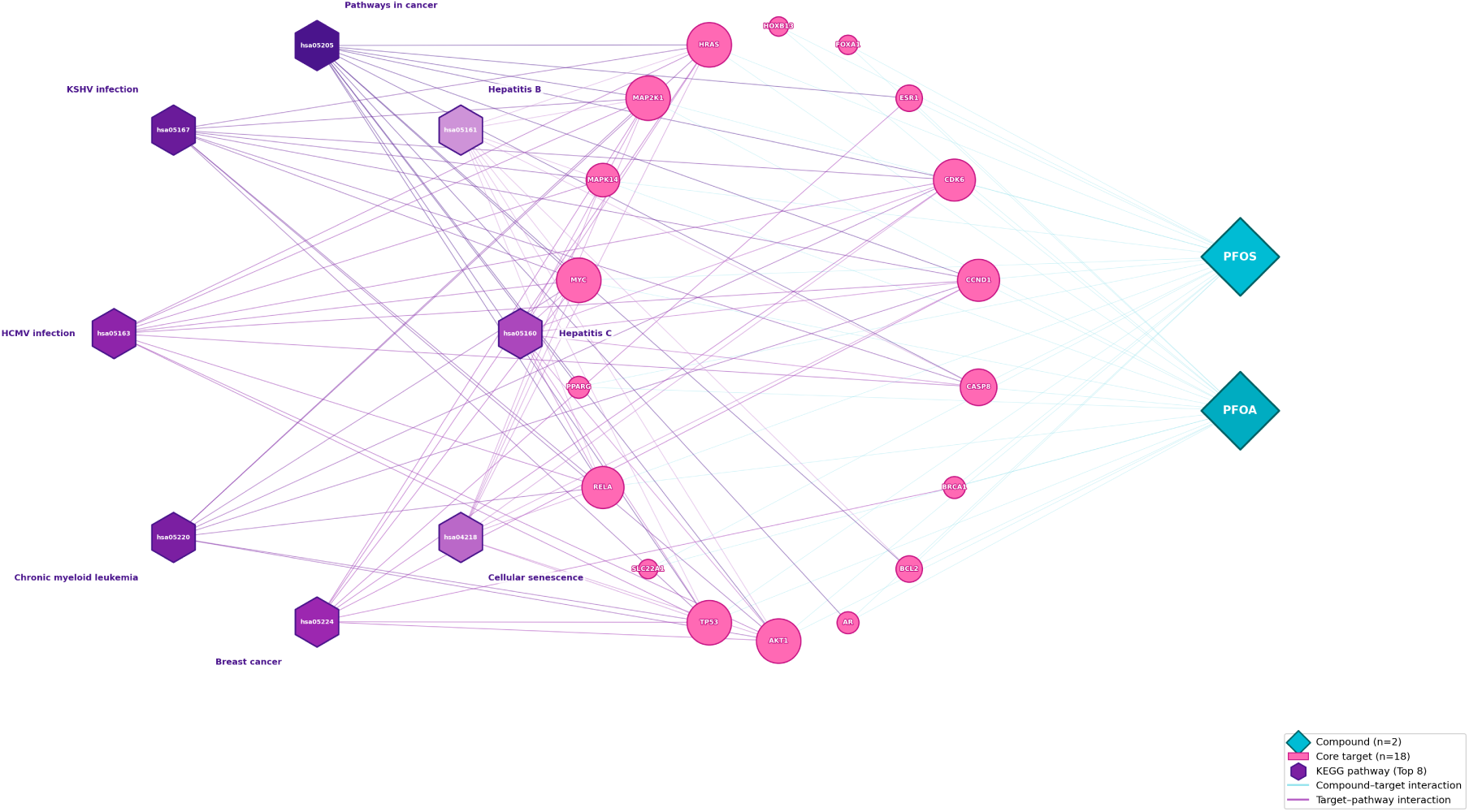
Three-layer compound-target-pathway network diagram.

### 3.11 Molecular docking: strong binding of PFOS/PFOA to AR, AKT1, and PTEN

To verify the direct regulatory effects of PFOS/PFOA on core targets at the molecular interaction level, we performed molecular docking. Target selection for docking followed three criteria: (1) Prognostic significance in survival analysis: PTEN (TCGA-PRAD DFS padj < 0.05, significant prognostic association) and AKT1 (HR = 2.97, suggestive risk trend) exhibited the strongest prognostic associations in the clinical cohorts. (2) Machine learning consensus: AR (selected by 2/3 algorithms, supported by both RF and SVM, and additionally backed by GWAS genetic association evidence), AKT1 (3/3 consensus), and PTEN (3/3 consensus) were the targets with the strongest evidence in machine learning-based screening. (3) Biological prior knowledge and pathway importance: AR is a core driver of prostate carcinogenesis, and the AR signaling axis is a major therapeutic target in prostate cancer; PTEN loss is the most frequent genomic alteration in prostate cancer (∼40% of cases), and its phosphatase activity directly regulates the PI3K-AKT-mTOR axis; AKT1 is the central kinase of the PI3K-AKT pathway, integrating multiple upstream signals.

Together, these three targets encompass the two core mechanistic axes of PFOS/PFOA toxicity—the AR-mediated endocrine disruption axis and the PI3K-AKT proliferation axis. The results showed that both PFOS and PFOA exhibited strong binding affinities toward all three targets, with PFOS consistently demonstrating stronger binding than PFOA for each target (Figure 11A, D). The AR ligand-binding domain displayed the highest binding affinity (PFOS: −9.49 kcal/mol; PFOA: −8.56 kcal/mol), with the binding energy comparable to that of the natural ligand testosterone, suggesting that PFOS/PFOA may exert endocrine-disrupting effects through competitive binding to AR (Figure 11B, E). The affinity distribution of the nine docking conformations indicated that the AR–PFOS docking conformations were the most clustered, reflecting a high degree of complementarity and stability of the binding pocket (Figure 11C). Detailed docking parameters are presented in Table 2^[16,17]^.

**Figure 11.**
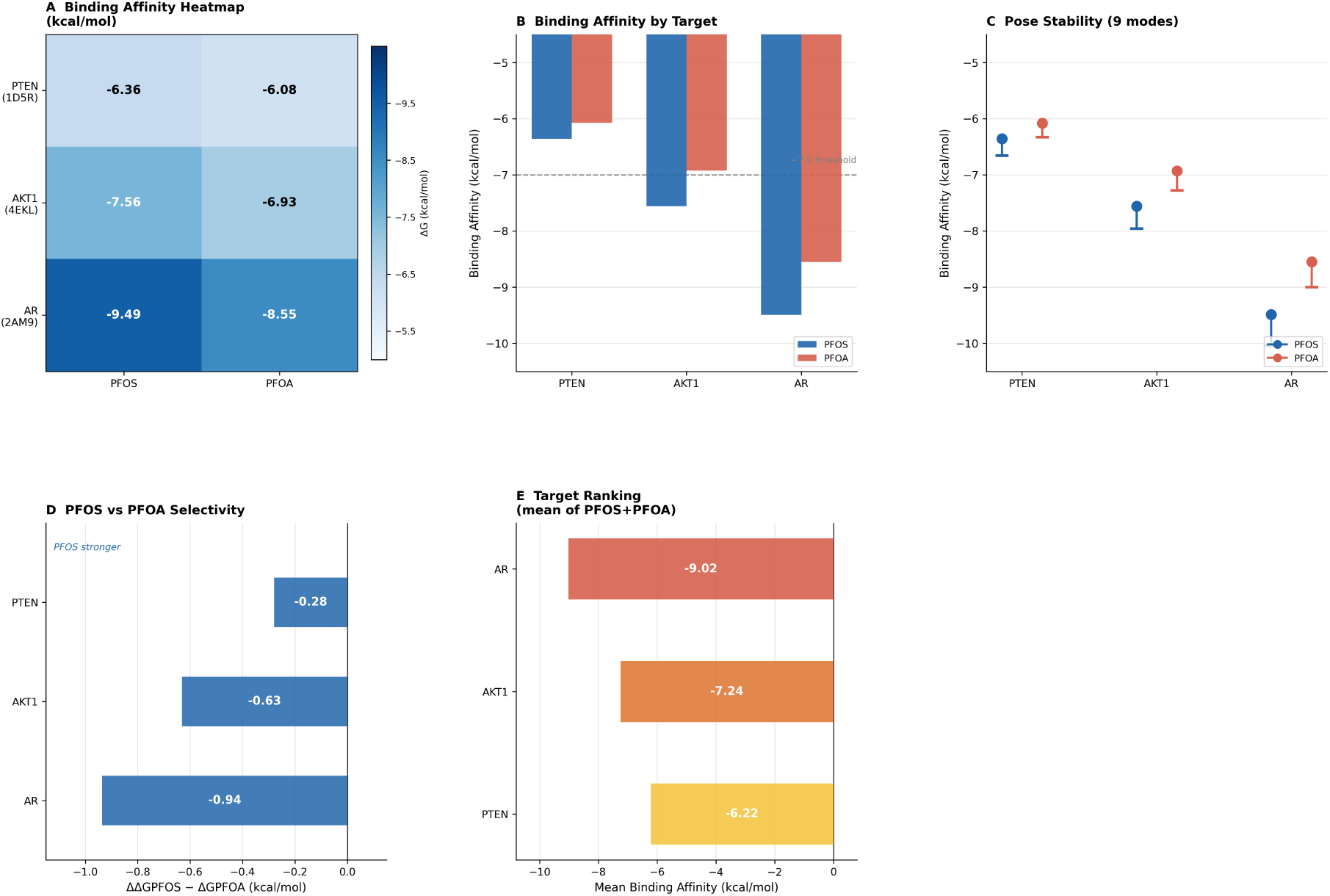
(A) Heatmap of binding affinity.(B) Grouped bar plot of binding affinity for each target.(C) Lollipop plot of docking conformation stability.(D) Plot of binding selectivity difference between PFOS and PFOA.(E) Ranking plot of mean binding affinity for each target.

**Table 2.** Summary of molecular docking between PFOS/PFOA and AR, AKT1 and PTEN.

| Target | PDB ID | Resolution | Binding Pocket | PFOS $\Delta G$ (kcal/mol) | PFOA $\Delta G$ (kcal/mol) | Rating |
| --- | --- | --- | --- | --- | --- | --- |
| AR | 2AM9 | 1.64 Å | Ligand-binding domain (LBD) | -9.491 | -8.555 |  |
| AKT1 | 4EKL | 2.00 Å | ATP-binding site | -7.559 | -6.928 |  |
| PTEN | 1D5R | 2.10 Å | Phosphatase active site | -6.356 | -6.077 |  |

### 3.12 PFAS-PTS risk stratification and survival validation

Based on the univariate Cox regression coefficients of the 18 core targets, we constructed a PFAS-PTS risk score purely weighted by Cox coefficients, with the formula: PFAS-PTS = Σ(βᵢ × Expression_geneᵢ). Patients were stratified into high- and low-risk groups using the median score of the GSE16560 cohort as the cutoff. Survival analysis demonstrated that in the GSE16560 training cohort, patients in the high-risk group had significantly shorter overall survival than those in the low-risk group (log-rank p=8.4×10−6p=8.4×10−6, HR = 1.55, 95% CI: 1.33–1.79) (Figure 12A). In the TCGA-PRAD independent validation cohort, the high-risk group exhibited a significantly elevated risk of biochemical recurrence (log-rank p=2.7×10−3p=2.7×10−3, HR = 1.45, 95% CI: 1.18–1.78) (Figure 12B), confirming that the PFAS-PTS score possesses robust cross-cohort prognostic stratification capability.

**Figure 12.**
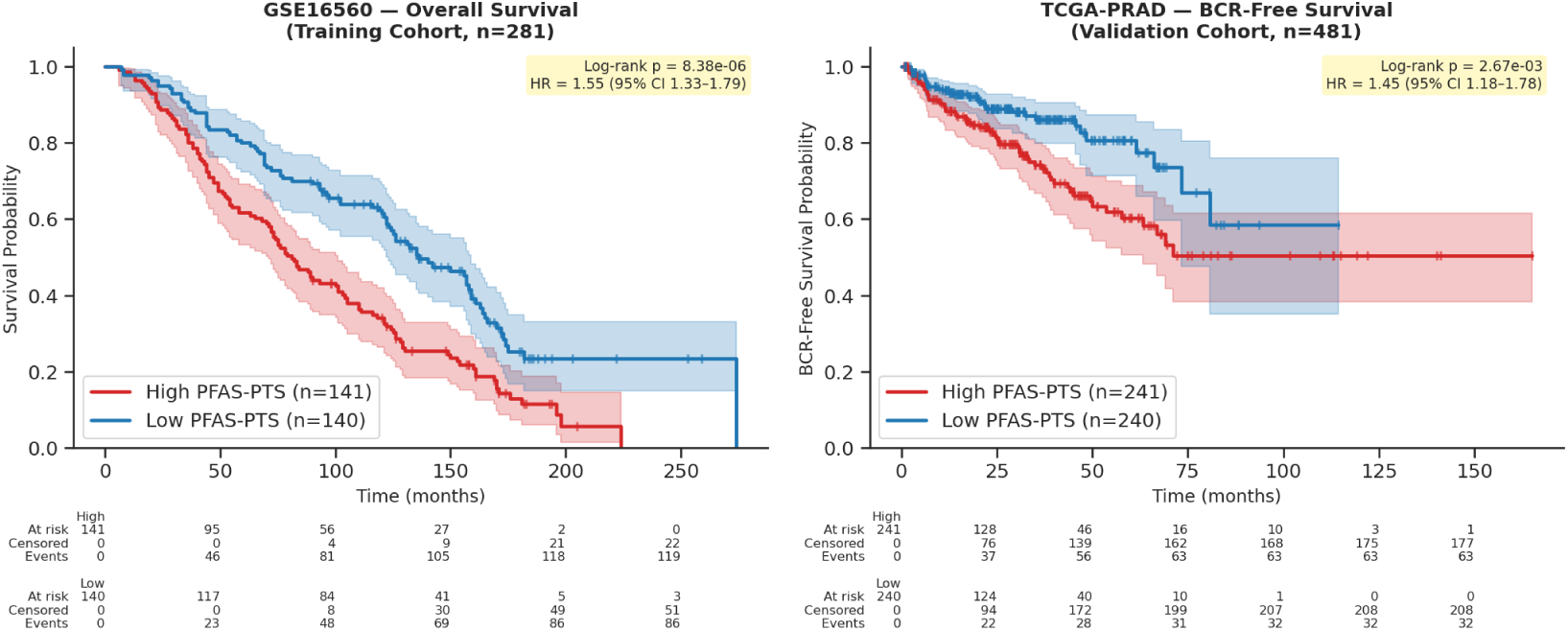
(A) Kaplan-Meier (KM) curve of overall survival for the GSE16560 cohort.(B) Kaplan-Meier (KM) curve of biochemical recurrence-free survival for the TCGA-PRAD cohort.

### 3.13 GSVA pathway analysis: upregulation of G2M/E2F and downregulation of androgen response

To elucidate the biological basis underlying the prognostic stratification capacity of the PFAS-PTS score, we performed single-sample gene set enrichment analysis (ssGSEA) to compare the activity of core oncogenic pathways between the high- and low-risk groups. The results demonstrated that in both the GSE16560 and TCGA-PRAD independent cohorts, the G2M checkpoint and E2F target gene pathways were significantly upregulated in the high-risk group (padj < 0.001), indicating pronounced aberrant cell cycle activation and upregulation of proliferative programs in high-risk patients. The PI3K/AKT/mTOR signaling pathway was significantly enriched in the high-risk group of the TCGA-PRAD cohort (padj = 2.1 × 10⁻⁴), consistent with the mechanistic hypothesis that PFAS activate the PI3K-AKT pathway. The androgen response pathway was significantly downregulated in the high-risk group of the TCGA-PRAD cohort (padj = 4.8 × 10⁻⁴), suggesting that high-risk patients may progress toward an androgen-independent phenotype. Of the seven pathways examined, six exhibited concordant directional changes across both cohorts, demonstrating favorable cross-cohort reproducibility (Figure 13).

**Figure 13.**
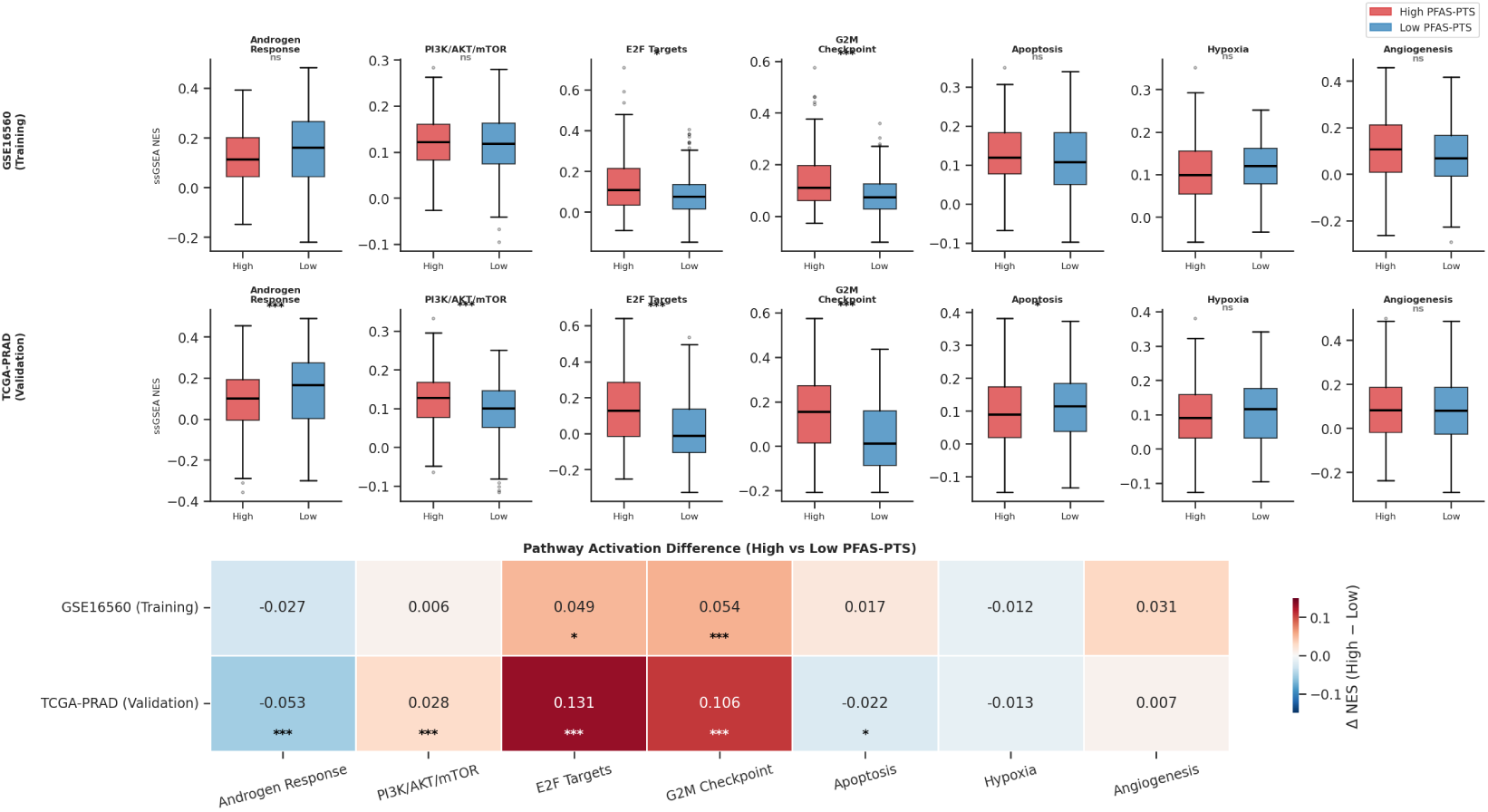
Box plots of pathway ssGSEA scores and heatmap of effect sizes.

### 3.14 TME deconvolution of high- and low-risk groups

We constructed a prostate cancer-specific signature matrix based on single-cell transcriptomic data from GSE141445 and performed TME deconvolution using NNLS to resolve differences in cellular composition between the PFAS-PTS high- and low-risk groups. In the TCGA-PRAD cohort, the high-risk group exhibited a significantly higher proportion of tumor epithelial cells (median 0.379 vs. 0.358, p=2.4×10−4p=2.4×10−4) and a significantly lower proportion of CD8⁺ cytotoxic T cells (median 0.085 vs. 0.098, p=5.7×10−4p=5.7×10−4) compared with the low-risk group, accompanied by increasing trends in the proportions of regulatory T cells and M2 tumor-associated macrophages (Figure 14). The GSE16560 cohort displayed completely concordant trends, confirming that patients with high PFAS-PTS scores present typical features of an immunosuppressive TME. These findings suggest that PFAS exposure may promote immune evasion and progression of prostate cancer by remodeling the TME^[22,23]^.

**Figure 14.**
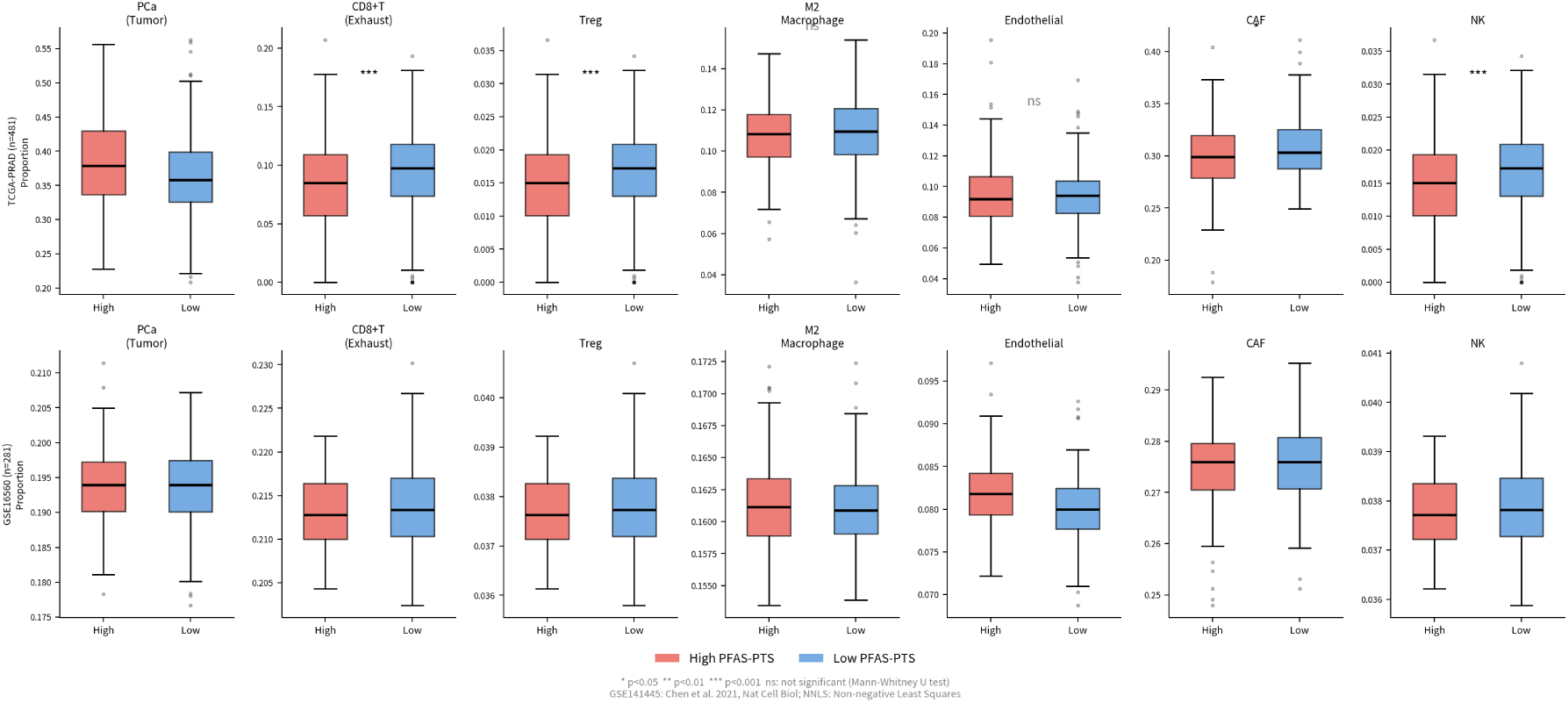
(A) Box plots of cell proportions for the TCGA-PRAD cohort.(B) Box plots of cell proportions for the GSE16560 cohort.

### 3.15 ODE kinetic simulation

To transcend the limitations of static omics analyses and quantitatively delineate the dynamic regulatory effects of PFAS exposure on the prostate cancer TME, we constructed a 10-dimensional ODE kinetic system incorporating six cell types and four core microenvironmental variables. Initial conditions for the model were derived from the NNLS deconvolution results of the TCGA-PRAD cohort, and the PFAS exposure effect was modeled using a Hill function. Dose-gradient simulations revealed that as the PFAS concentration increased, the tumor epithelial cell (PCa) proportion rose in a dose-dependent manner, whereas the CD8⁺ T cell proportion decreased dose-dependently; concomitantly, immunosuppression-associated indicators, including Tregs, M2 macrophages, lactate, and IL-6, all exhibited dose-dependent elevations, establishing a clear exposure-dose–biological-effect relationship (Figure 15A). Under unexposed conditions (0 ng/mL), the immune system effectively controlled tumor proliferation; however, at 20 ng/mL PFAS, the tumor-proliferative effect exceeded the immune killing capacity, resulting in a net proliferative state. Comparison of kinetic trajectories between the high and low PFAS-PTS groups demonstrated that the high-risk group possessed stronger tumor proliferative capacity and more pronounced immunosuppressive effects, fully consistent with the static omics findings (Figure 15B). Monte Carlo uncertainty analysis (n = 1,000, with ±20% uniform perturbations applied to all 47 kinetic parameters) demonstrated that the qualitative conclusions of the model—specifically, the directionality of the PFAS dose–effect relationship—remained highly robust within the parameter uncertainty range, with 100% of simulations converging successfully. OAT parameter sensitivity analysis (±20% perturbation) identified the PCa proliferation rate (r_PCa), the carrying capacity (K), and the maximum PFAS effect (E_max) as the parameters exerting the greatest influence on the model output (Figure 16)^[24]^.

**Figure 15.**
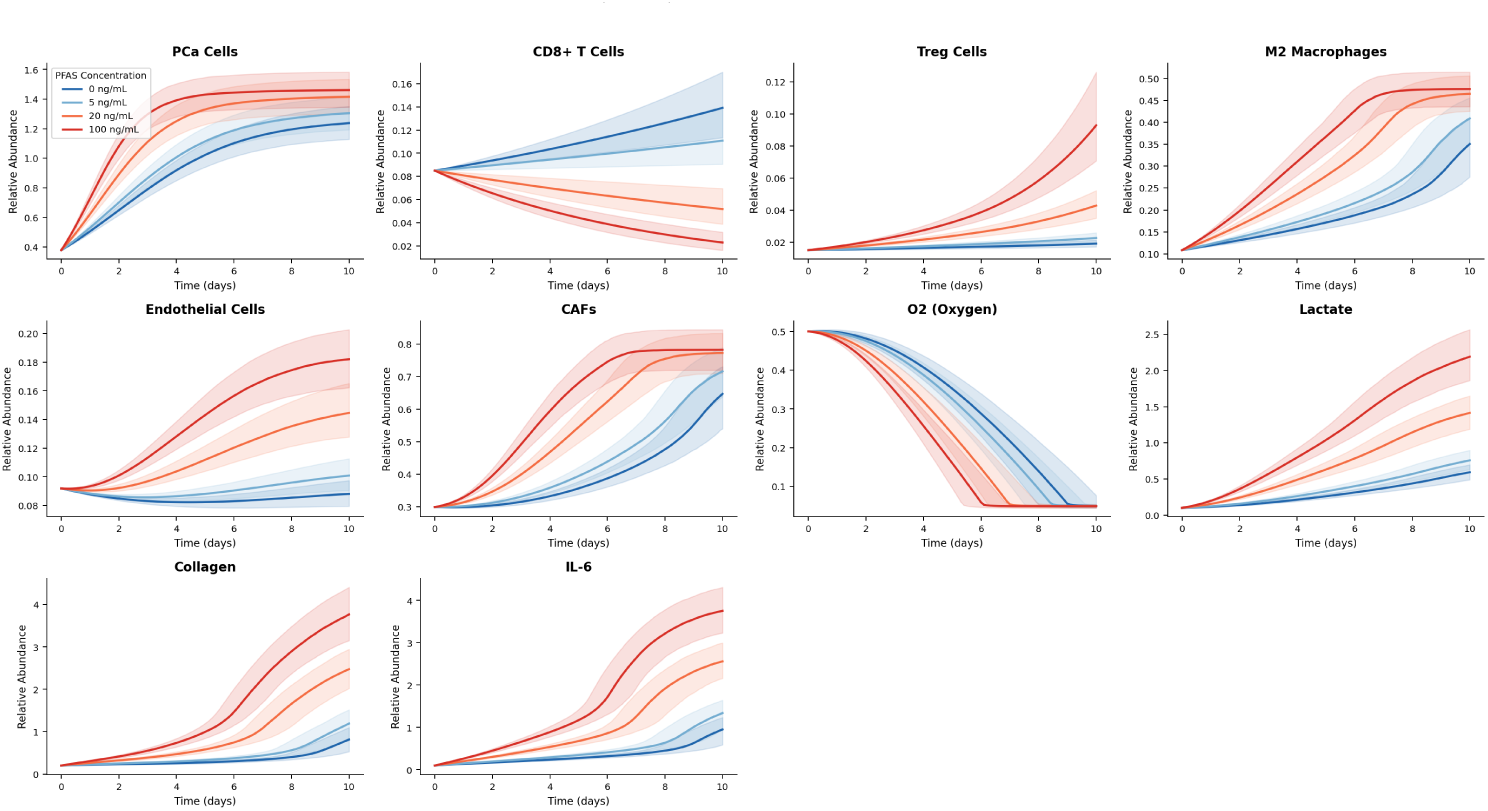
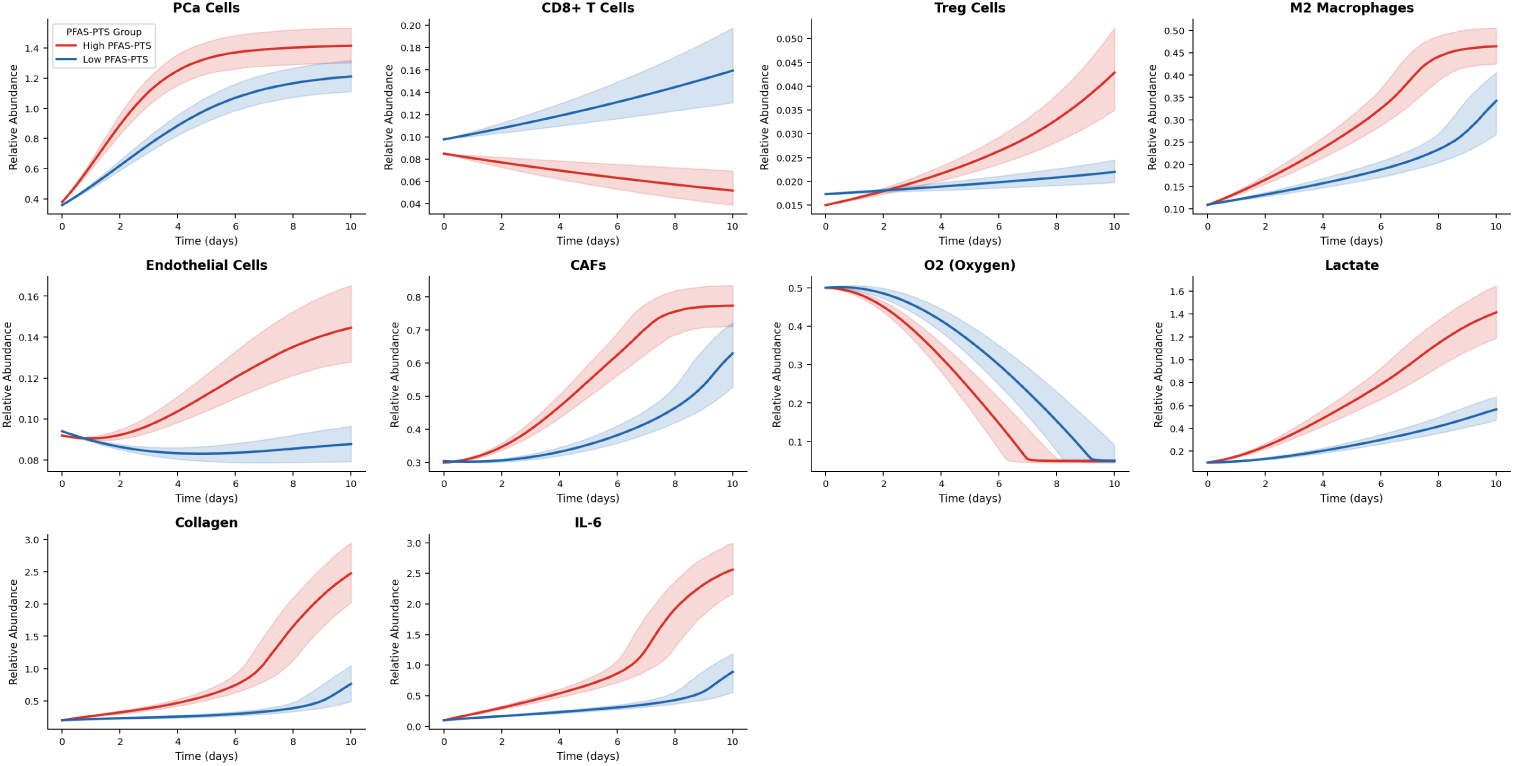
(A) Dynamic trajectory plots of state variables across PFAS dose gradients.(B) Comparative dynamic trajectory plots of the TME between high and low PFAS-PTS groups.

**Figure 16.**
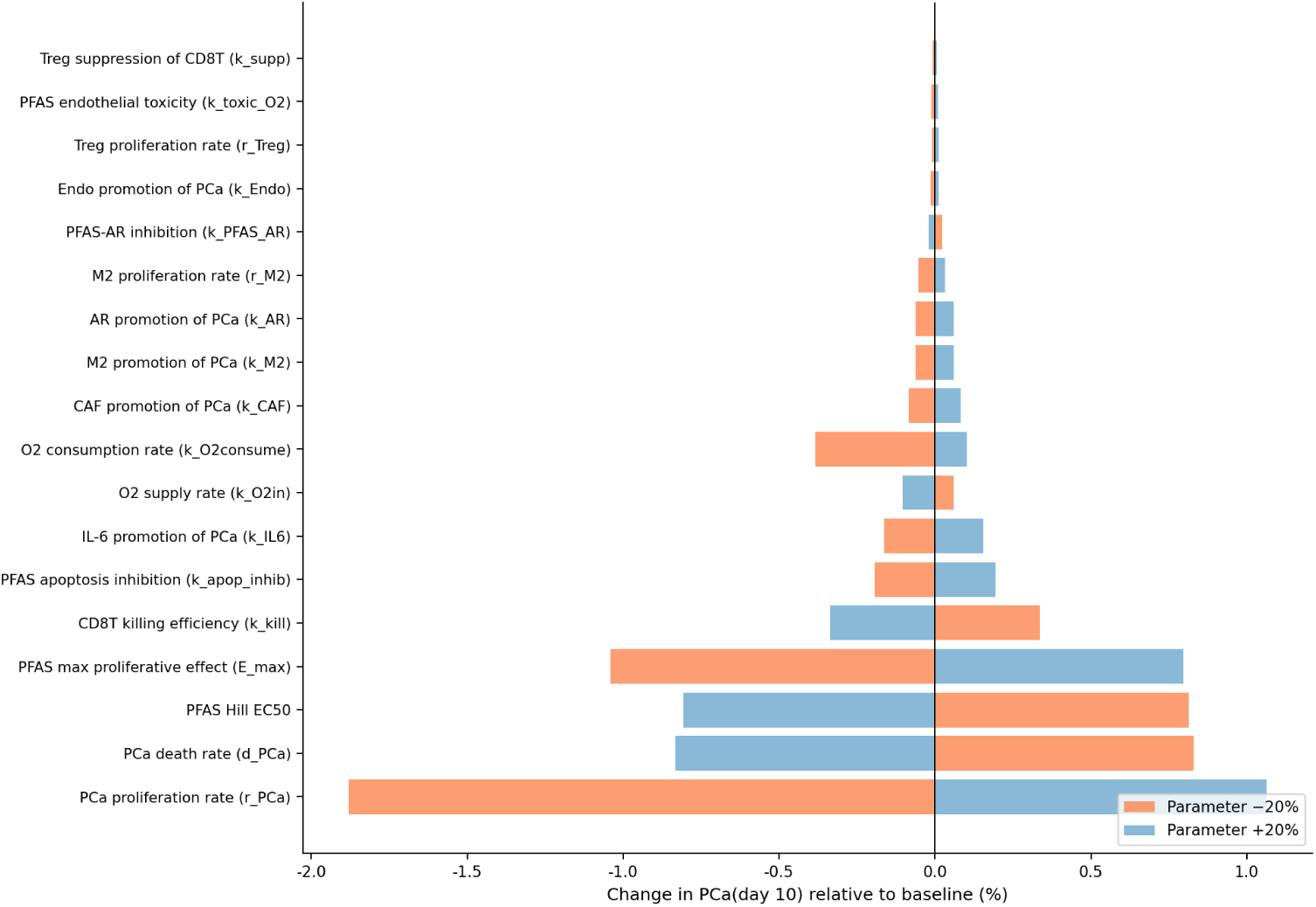
Tornado plot of parameter sensitivity analysis for the ODE model.

### 3.16 Drug intervention prediction

Based on the constructed ODE model, we systematically evaluated the inhibitory efficacy of five clinical therapeutic strategies against prostate cancer under a background of high PFAS exposure. Under high-risk initial conditions with 20 ng/mL PFAS exposure, all intervention strategies reduced the tumor cell proportion to varying degrees. Among them, enzalutamide combined with Alpelisib (dual AR and PI3K inhibition) achieved the optimal tumor inhibitory effect, with an inhibition rate of 33.9%; palbociclib (CDK4/6 inhibitor) monotherapy ranked second (18.0%); enzalutamide monotherapy yielded an inhibition rate of 16.2%; and Alpelisib monotherapy and venetoclax combined with bevacizumab exhibited comparatively limited inhibitory effects (Figure 17). The model also revealed that PFAS-induced apoptosis resistance can attenuate the therapeutic efficacy of CDK4/6 inhibitors, suggesting that combination with an apoptosis restoration strategy may further enhance treatment outcomes.

**Figure 17.**
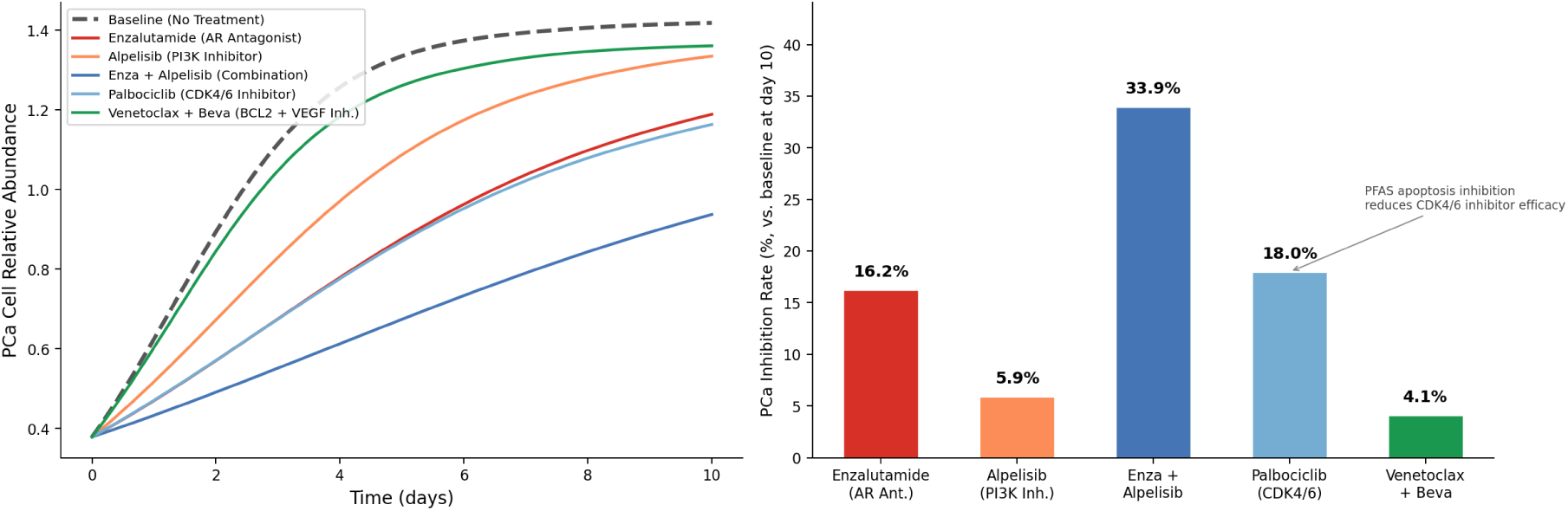
(A) Dynamic trajectory plot of PCa cell abundance.(B) Bar plot of inhibition rate of each drug regimen.

### 3.17 Global sensitivity and uncertainty quantification analysis of the ODE model

Morris global screening systematically assessed 52 parameters of the ODE system and revealed that the PCa proliferation rate (r_PCa), tumor carrying capacity (K), CD8⁺ T cell proliferation rate, CD8⁺ T cell killing efficiency (k_kill), and the maximum PFAS effect intensity (E_max) are high-impact parameters (μ*>0.02), results that are highly consistent with the OAT sensitivity analysis (Figure 18). Sobol global sensitivity analysis further quantified the contribution proportions of the key parameters, demonstrating that r_PCa exerts the most dominant effect on the model output; both its first-order and total-effect indices were significantly higher than those of the other parameters, and the interaction effects among parameters were extremely weak (ΣS_T≈1.0), suggesting that model behavior is primarily driven linearly by core proliferation and toxicity parameters (Figure 19).

**Figure 18.**
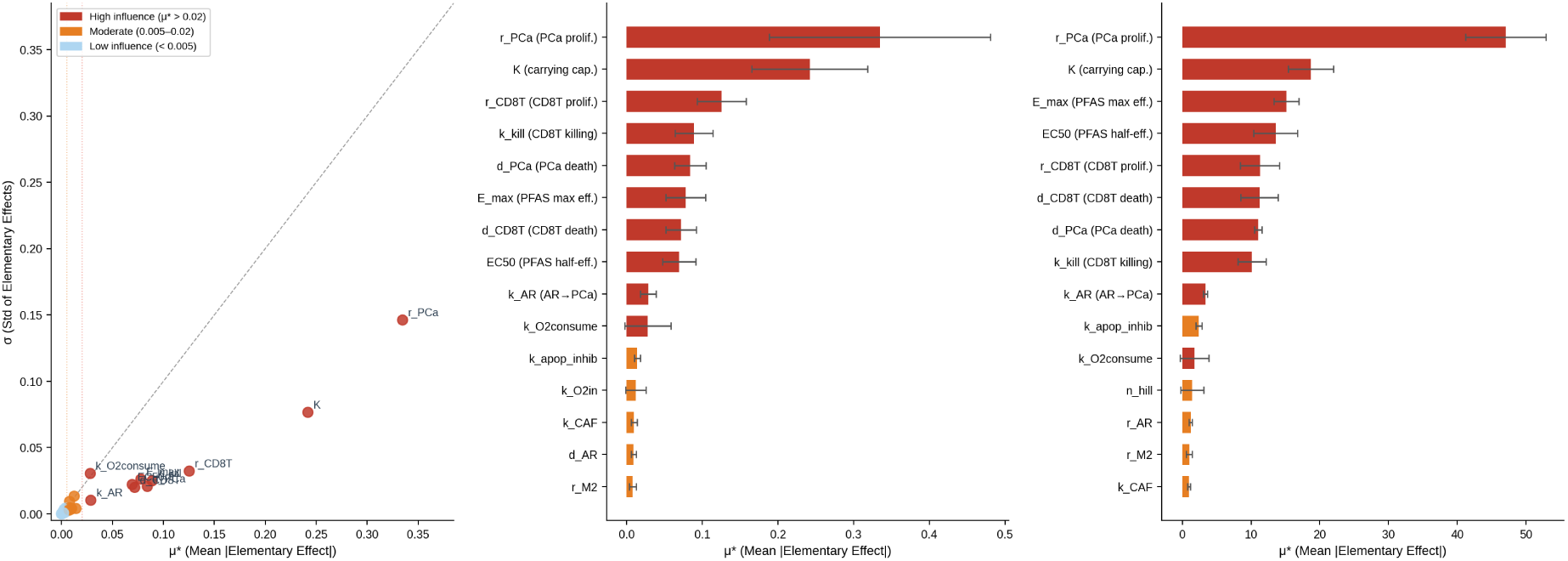
(A) μ*-σ scatter plot of Morris global sensitivity analysis.(B) Ranking plot of parameter importance for PFAS dose-response effect.(C) Ranking plot of parameter importance for drug intervention effect.

**Figure 19.**
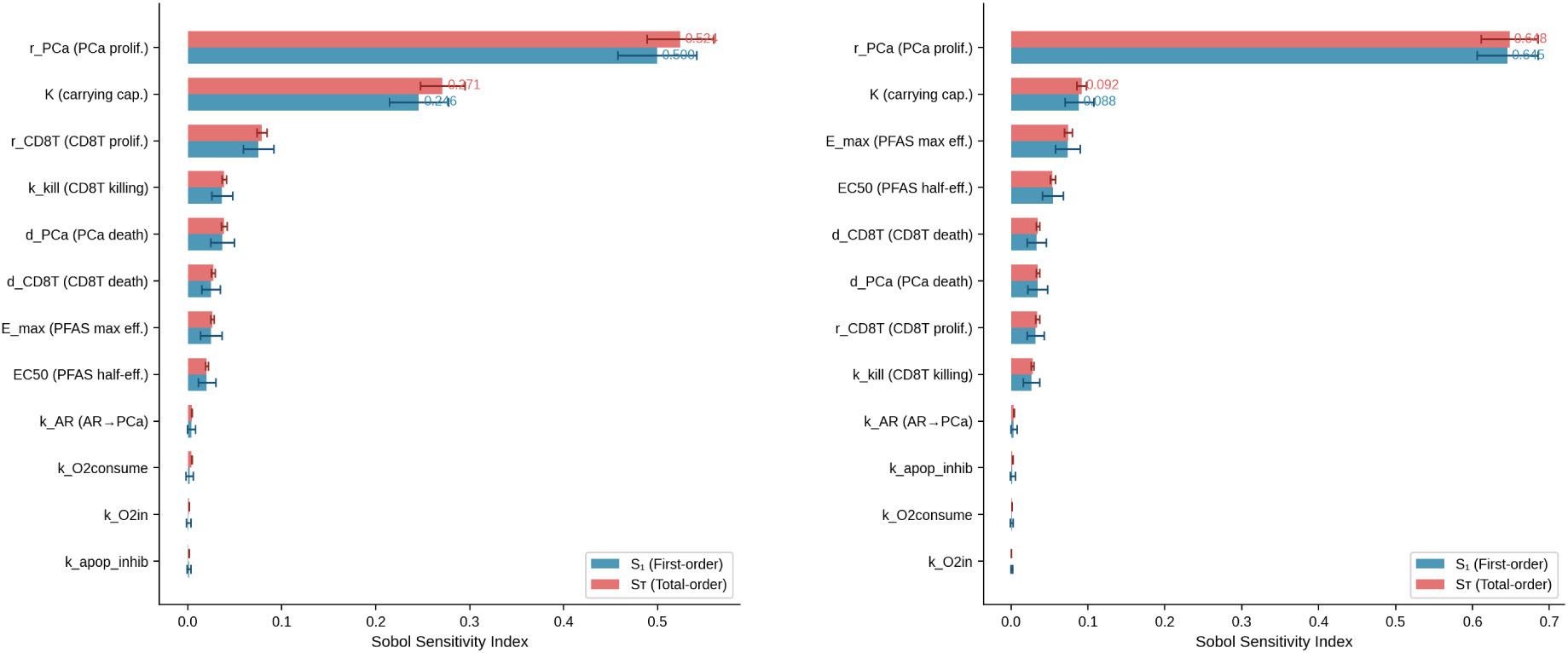
(A) Sobol sensitivity index plot for PFAS dose-response effect.(B) Sobol sensitivity index plot for drug intervention effect.

Monte Carlo parameter perturbation analysis (n = 2,000, ±20% uniform perturbations) confirmed that all drug intervention strategies maintained stable tumor-inhibitory effects within the range of parameter uncertainty. Notably, the combination of enzalutamide and Alpelisib retained a median inhibition rate of 33.9% with a robust 95% confidence interval, significantly outperforming other monotherapies or combination regimens, indicating that the superiority of this combination strategy is independent of specific parameter values and possesses reliable predictive stability (Figure 20). These results validate the scientific robustness of the ODE system constructed in this study from a modeling perspective and provide quantitative support for the computational prediction of intervention strategies for PFAS-associated prostate cancer.

**Figure 20.**
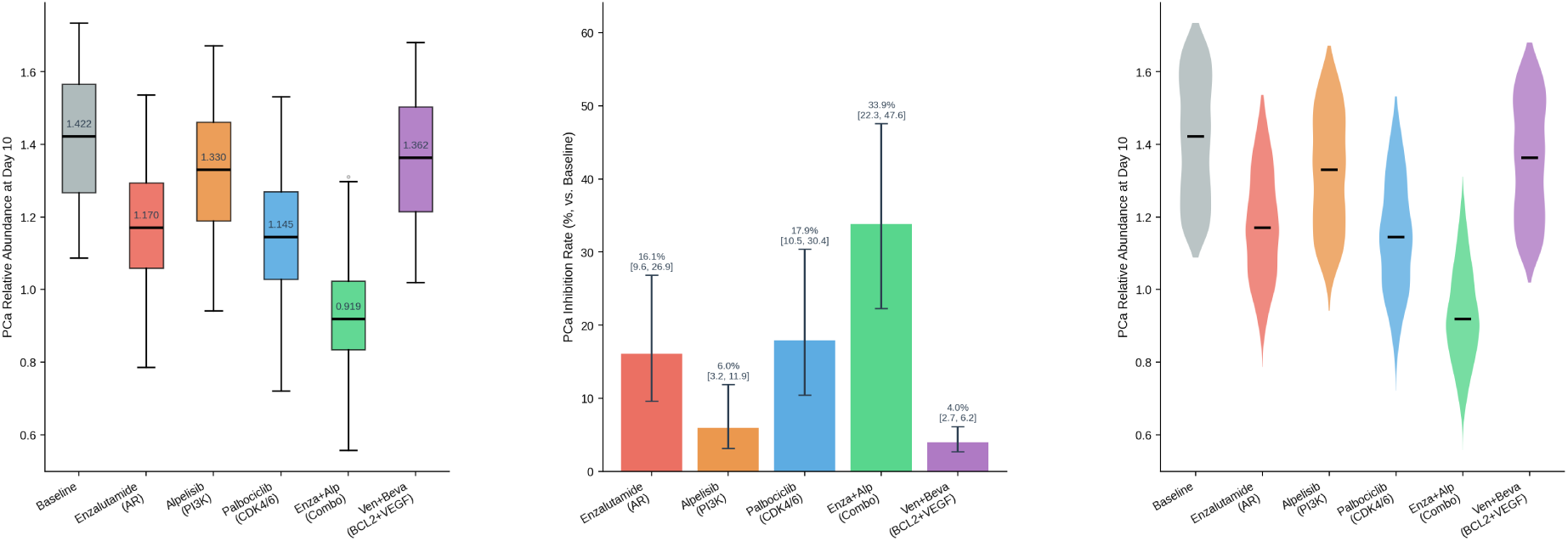
(A) Box plots of PCa cell abundance distribution.(B) Bar plot of drug inhibition rates with 95% confidence intervals.(C) Violin plot of inhibition rate distribution.

## 4 Discussion

As the second most common malignancy in men worldwide, PCa has exhibited a steadily rising incidence in industrialized countries, and exposure to environmental pollutants has been established as an important risk factor. PFOS and PFOA, two of the most representative persistent organic pollutants, have been supported by multiple cohort studies for their epidemiological association with increased PCa risk. However, the precise molecular mechanisms by which combined PFOS/PFOA exposure promotes prostate carcinogenesis and progression—particularly the dynamic remodeling of the TME—have long remained largely unexplored in a systematic and quantitative manner. Traditional single-target approaches are inherently inadequate to capture the characteristic “multi-target, multi-pathway, network-based” toxicity of PFAS, while existing network pharmacology studies are largely confined to static target–pathway association analyses and are unable to reveal the dynamic evolutionary patterns of tumor biological behavior under environmental exposure. In this study, we innovatively constructed an integrated analytical framework that combines nine modules of network pharmacology and computational biology, and for the first time achieved a comprehensive chain-based systematic deduction from “molecular target identification → pathway dysregulation dissecting → TME remodeling → kinetic simulation → drug intervention prediction”. Through multi-database cross-validation, machine learning-based independent screening, GWAS genetic association corroboration, and dual-cohort clinical survival analysis, we precisely identified 18 PFOS/PFOA–PCa core targets. Further, by integrating PCa-specific TME deconvolution and a 10-dimensional ODE kinetic simulation, we quantitatively elucidated the dose–effect relationship by which PFAS exposure drives PCa progression and predicted an optimal combination therapeutic strategy. This study not only provides a novel theoretical perspective on the molecular mechanisms of PFAS-associated PCa but also offers translatable computational evidence for its risk assessment and precision therapy ^[1,3,25]^.

At the molecular mechanistic level, we first focused on the core driver pathway of prostate carcinogenesis and progression—the AR signaling axis. A focused AR axis analysis revealed that 61% of the 100 common targets harbor functional associations with the AR signaling network, covering eight key functional modules including ligand binding, co-regulator recruitment, kinase crosstalk, downstream transcriptional activation, and cell cycle regulation. This finding strongly supports the central hypothesis that PFAS act as endocrine-disrupting chemicals. In contrast to earlier studies that generally considered PFAS to only exhibit weak antagonistic effects through competitive binding to the AR ligand-binding domain (LBD), our study demonstrated that PFOS/PFOA can simultaneously target multiple nodes of the AR signaling pathway. On the one hand, molecular docking results confirmed that PFOS (−9.49 kcal/mol) and PFOA (−8.56 kcal/mol) possess binding affinities for the AR-LBD comparable to that of the natural ligand testosterone, likely forming hydrogen bonds with key residues such as Arg752 and Asn705 to competitively block androgen–AR binding. On the other hand, PFAS may also regulate AR phosphorylation and nuclear translocation by targeting kinases such as AKT1 and PTEN, thereby affecting its transcriptional activity. This “multi-site synergistic interference” model explains why PFAS exhibit only weak anti-androgenic activity in vitro yet can significantly promote PCa progression in vivo ^[4,20]^.

Beyond directly interfering with the AR signaling network, PFOS/PFOA also cooperatively promote prostate cancer progression through aberrant activation of the PI3K-AKT signaling pathway. Aberrant activation of the PI3K-AKT pathway is a key mechanism underlying castration-resistant progression; PTEN loss or PI3K-AKT pathway activation occurs in approximately 40% of primary PCa and 70% of castration-resistant prostate cancer (CRPC). Among the 18 core targets identified in this study, AKT1, PTEN, and MTOR are key molecules of the PI3K-AKT pathway, and focused enrichment analysis showed that this pathway is significantly upregulated in the PFAS-PTS high-risk group. Molecular docking further confirmed that PFOS/PFOA can directly bind to the AKT1 kinase domain (PFOS: −7.56 kcal/mol; PFOA: −6.93 kcal/mol) and inhibit PTEN phosphatase activity (PFOS: −6.36 kcal/mol; PFOA: −6.08 kcal/mol), thereby releasing PTEN-mediated negative regulation of the PI3K-AKT pathway and leading to its sustained activation. Notably, there exists a complex negative feedback regulatory mechanism between AR and PI3K-AKT pathways: AR activation can inhibit the PI3K-AKT pathway by upregulating PTEN expression, whereas PI3K-AKT pathway activation can promote androgen-independent AR activation through phosphorylation of AR and its co-regulators. Our study revealed that PFAS can simultaneously disrupt these two core pathways, creating a vicious cycle of “AR signaling disorder–PI3K-AKT hyperactivation”, which may represent an important molecular basis for PFAS exposure-accelerated progression toward castration resistance ^[26,27]^.

The synergistic activation of AR and PI3K-AKT pathways ultimately converges on the deregulation of cell cycle control, which is a hallmark of cancer cells acquiring unlimited proliferative capacity. Our GSVA pathway analysis showed that both the G2M checkpoint and E2F target gene pathways were significantly upregulated in the PFAS-PTS high-risk group in two independent cohorts (padj < 0.001), indicating that PFAS exposure promotes tumor cell proliferation primarily by driving the G2/M transition of the cell cycle. E2F transcription factors are key regulators of both G1/S and G2/M transitions and represent common downstream targets of the AR and PI3K-AKT pathways. Through activation of AR and PI3K-AKT pathways, PFAS may upregulate the expression of cyclins such as CCND1 and CDK6, promote Rb protein phosphorylation, and release E2F, thereby triggering cell cycle progression. In addition, our dual-cohort survival analysis showed that RELA was significantly associated with poor prognosis in both cohorts, suggesting that activation of the NF-κB signaling pathway may also contribute to PFAS-induced PCa progression. NF-κB not only directly regulates the expression of cell cycle and apoptosis-related genes but also engages in crosstalk with AR and PI3K-AKT pathways, further amplifying the carcinogenic effects of PFAS ^[18]^.

Notably, these carcinogenic effects of PFAS do not occur independently of genetic background but instead exhibit significant interaction with the genetic susceptibility of prostate cancer. As validated through the GWAS Catalog, 13 of the 18 core targets possessed genome-wide significant genetic associations with PCa (p < 5 × 10⁻⁸), among which HOXB13 (p = 6 × 10⁻³⁴), KLK3 (p = 2 × 10⁻²⁸), and NKX3-1 (p < 10⁻²⁰) are well-established strong PCa susceptibility genes. This finding indicates that PFAS may amplify an individual’s genetic risk for PCa by regulating the expression or function of genetic susceptibility genes. For example, HOXB13 is a key transcription factor in prostate development and tumorigenesis, and its G84E mutation significantly increases the risk of PCa onset; our study identified HOXB13 as the strongest positive risk factor in the PFAS-PTS score (β = +0.239), suggesting that PFAS exposure may further potentiate its oncogenic effect by upregulating HOXB13 expression. This “environmental exposure–genetic susceptibility” interplay provides an important theoretical basis for precision prevention of PCa.

In addition to directly regulating the intrinsic proliferative signaling of tumor cells, PFAS exposure indirectly promotes PCa progression by remodeling the TME. The TME serves as the “soil” for tumor cell survival and progression, and the formation of an immunosuppressive microenvironment is a major cause of tumor immune evasion and therapeutic resistance. Previous studies on PFAS and tumor immunity have predominantly focused on systemic immunotoxicity, whereas investigations into how PFAS exposure reshapes the local solid tumor microenvironment remain very limited. In this study, using a specific signature matrix constructed from GSE141445 PCa single-cell transcriptomic data and applying NNLS deconvolution to the TCGA-PRAD and GSE16560 cohorts, we systematically revealed, for the first time, the TME characteristics of PFAS-PTS high-risk PCa patients: a significantly increased proportion of tumor epithelial cells, decreased CD8⁺ cytotoxic T cell infiltration, and elevated proportions of regulatory T cells (Tregs) and M2 tumor-associated macrophages (M2-TAMs). These features are highly consistent with an immune “cold tumor” phenotype, suggesting that PFAS exposure can promote immune evasion of PCa by reshaping an immunosuppressive microenvironment ^[22,23,28]^.

To overcome the limitations of static deconvolution analysis and quantitatively characterize the dynamic regulatory patterns of PFAS exposure on individual TME components, we constructed a 10-dimensional ODE kinetic model incorporating six cell types and four core microenvironmental variables. Simulation results demonstrated that PFAS exposure promotes tumor cell proliferation in a dose-dependent manner while simultaneously suppressing the anti-tumor immune response of CD8⁺ T cells and inducing the recruitment and activation of Tregs and M2-TAMs. Under PFAS-free conditions (0 ng/mL), the immune system effectively controlled tumor cell proliferation; however, when the PFAS concentration reached 20 ng/mL (a reference concentration for high-exposure groups in epidemiological settings), PFAS-driven tumor proliferative effects exceeded the immune killing capacity, resulting in a net proliferative state. Monte Carlo uncertainty analysis (n = 1,000 parameter perturbations) confirmed that these qualitative conclusions are highly robust within the range of parameter uncertainty ^[24]^.

Based on the above static and dynamic analytical results, the molecular mechanisms by which PFAS induce an immunosuppressive microenvironment can be summarized at three levels. First, PFAS can directly act on immune cells, inhibiting the proliferation and cytotoxic function of CD8⁺ T cells while promoting the differentiation and immunosuppressive activity of Tregs. Second, PFAS-activated PI3K-AKT and NF-κB pathways can induce tumor cells to secrete cytokines such as IL-6 and TGF-β, which further recruit and activate M2-TAMs and Tregs, forming an immunosuppressive positive feedback loop. Furthermore, PFAS can also hinder immune cell infiltration into tumor tissue by promoting angiogenesis and extracellular matrix remodeling. Our ODE model incorporated microenvironmental variables including O₂ concentration, lactate, collagen, and IL-6, and simulations showed that PFAS exposure leads to exacerbated tumor hypoxia, lactate accumulation, and collagen deposition—microenvironmental alterations that not only directly suppress immune cell function but also promote tumor cell invasion and metastasis.

These mechanistic discoveries provide clear target directions for the precision treatment of PFAS-associated PCa. Accordingly, this study systematically evaluated the potential efficacy of multiple clinically used therapeutic regimens through ODE kinetic simulation. Currently, the treatment of PCa primarily includes endocrine therapy, chemotherapy, targeted therapy, and immunotherapy, but for CRPC, the efficacy of existing regimens remains quite limited. PFAS exposure not only promotes PCa progression but may also influence tumor cell sensitivity to therapeutic drugs, making the development of specific treatment strategies for PFAS-associated PCa of significant clinical importance. Simulation results showed that the combination of enzalutamide and Alpelisib (dual inhibition of AR and PI3K) achieved the optimal tumor cell inhibition (33.9%), significantly outperforming monotherapies ^[29,30]^.

This result is highly consistent with the negative feedback regulatory mechanism between the AR and PI3K-AKT pathways. Previous studies have shown that the use of AR antagonists alone (e.g., enzalutamide) can lead to compensatory activation of the PI3K-AKT pathway, thereby causing resistance, whereas PI3K inhibitors alone (e.g., Alpelisib) can reprogram AR signaling by relieving its inhibition. Therefore, combined inhibition of AR and PI3K-AKT pathways can block this negative feedback regulation and exert a synergistic anti-tumor effect. The results of the IPATential150 phase III clinical trial showed that abiraterone combined with Ipatasertib (an AKT inhibitor) significantly prolonged progression-free survival in PTEN-deficient metastatic CRPC patients, providing important clinical evidence that supports our computational predictions. This study further demonstrates that the advantage of such a combination strategy is even more pronounced under a background of high PFAS exposure, because PFAS can simultaneously activate both the AR and PI3K-AKT pathways, and monotherapy is insufficient to fully block their carcinogenic effects ^[27]^.

In addition to the advantage of the AR/PI3K dual-target combination, this study also revealed a phenomenon with important clinical implications: PFAS exposure can induce apoptosis resistance in tumor cells, thereby affecting the therapeutic efficacy of CDK4/6 inhibitors. GSVA pathway analysis showed that the apoptosis pathway was significantly downregulated in the PFAS-PTS high-risk group (TCGA-PRAD padj = 0.032), based on which we introduced an apoptosis inhibition parameter (k_apop_inhib = 0.378) into the ODE model. Simulation results indicated that the tumor inhibition rate of palbociclib monotherapy was only 18.0%, lower than expected; however, if combined with a BCL2 inhibitor (e.g., venetoclax) to restore the apoptotic pathway, efficacy could potentially be further enhanced. This finding explains why some PCa patients respond poorly to CDK4/6 inhibitors and suggests that in patients with high PFAS exposure, a combination strategy that restores apoptosis may be an effective approach to overcome drug resistance ^[31]^.

Compared with the above two strategies, venetoclax combined with bevacizumab exhibited the most limited efficacy, achieving only a 4.1% tumor cell inhibition rate. This may be because PFAS-driven PCa is primarily characterized by cell cycle dysregulation and enhanced proliferation, with apoptosis inhibition and angiogenesis serving merely as secondary mechanisms. This result suggests that treatment strategies for PFAS-associated PCa should preferentially target its core driver pathways (AR and PI3K-AKT) rather than secondary mechanisms.

The PFAS-PTS score constructed based on the core targets provides an actionable stratification tool for the clinical application of the aforementioned therapeutic strategies. This score can be used not only for prognostic stratification in PCa but also to guide treatment decision-making: for patients with a high PFAS-PTS, the combination regimen of enzalutamide plus Alpelisib should be prioritized, while for patients with a low PFAS-PTS score, endocrine therapy alone may prove sufficient. Furthermore, the PFAS-PTS score can serve as a biomarker for selecting patients most likely to benefit from combination therapy, thereby enabling precision treatment.

## 5 Conclusion

In this study, we constructed a multi-module analytical framework integrating network pharmacology and computational biology to systematically elucidate the molecular mechanisms by which combined PFOS/PFOA exposure promotes PCa. The study identified 100 shared PFOS/PFOA–PCa targets, from which 18 core targets were determined through multi-module screening, with significant enrichment in the AR signaling axis, the PI3K-AKT pathway, and cell cycle regulation. GSVA pathway analysis, TME deconvolution (based on the GSE141445 PCa-specific signature matrix), and ODE kinetic modeling collectively revealed, at multiple levels, the dynamic mechanisms by which PFAS exposure remodels the immunosuppressive TME. Drug intervention simulation suggested that enzalutamide combined with Alpelisib represents the most promising intervention strategy for PFAS-associated PCa (ODE model-predicted PCa inhibition rate: 33.9%).

## Consent

Not applicable.

## Acknowledge

This work was supported by awards from the Natural Science Foundation of Jiangsu Province (BK20231189), PanFeng Innovative Team Project of the The Third Affiliated Hospital of Soochow University(KY20252469) and Undergraduate Training Program for Innovation and Entrepreneurship,Soochow University(X2025102850485).

