## Supplementary Table S1 for "Systematic toxicological study of PFOS/PFOA co-exposure driving prostate cancer: Core target identification, TME immune remodeling, and combination drug prediction"

**Supplementary Table S1：**ODE Model Parameter Table

| **Parameter** | **Description** | **Value** | **Source** |
| --- | --- | --- | --- |
| r_PCa | PCa proliferation rate | 0.35 /day | Literature [32] |
| K_PCa | PCa carrying capacity | 1.0 (normalized) | Assumption |
| r_CD8T | CD8T proliferation rate | 0.20 /day | Literature [33] |
| K_CD8T | CD8T carrying capacity | 1.0 (normalized) | Assumption |
| r_Treg | Treg proliferation rate | 0.15 /day | Literature [33] |
| r_Endo | Endo proliferation rate | 0.10 /day | Literature [34] |
| r_M2 | M2 proliferation rate | 0.18 /day | Literature [33] |
| r_CAF | CAF proliferation rate | 0.12 /day | Literature [34] |
| α_M2_PCa | M2-mediated PCa promotion | 0.08 | Literature [33] |
| δ_CD8T_PCa | CD8T-mediated PCa killing | 0.15 | Literature [33] |
| α_PCa_Treg | PCa-induced Treg recruitment | 0.06 | Literature [33] |
| δ_Treg_CD8T | Treg-mediated CD8T suppression | 0.10 | Literature [33] |
| δ_M2_CD8T | M2-mediated CD8T suppression | 0.08 | Literature [33] |
| α_PCa_M2 | PCa-induced M2 polarization | 0.07 | Literature [33] |
| α_PCa_Endo | PCa-induced angiogenesis | 0.05 | Literature [34] |
| α_PCa_CAF | PCa-induced CAF activation | 0.06 | Literature [34] |
| β_PFAS | PFAS toxicity effect coefficient | 0.25 | Literature [35] |
| EC₅₀ | PFAS half-maximal concentration | 20 ng/mL | Literature [35] |

[32] Szklarczyk D, et al. (2023). The STRING database in 2023: protein-protein association networks and functional enrichment analyses for any of 12 967 organisms. Nucleic Acids Res, 51(D1), D638-D646. <https://doi.org/10.1093/nar/gkac1000>

[33] Cancer Genome Atlas Research Network. (2015). The molecular taxonomy of primary prostate cancer. Cell, 163(4), 1011-1025. <https://doi.org/10.1016/j.cell.2015.10.025>

[34] Hu J, Szymczak S. A review on longitudinal data analysis with random forest. Brief Bioinform. 2023;24(2):bbad002. doi:10.1093/bib/bbad002

[35] Trott O, Olson AJ. (2010). AutoDock Vina: improving the speed and accuracy of docking with a new scoring function, efficient optimization, and multithreading. J Comput Chem, 31(2), 455-461. https://doi.org/10.1002/jcc.21334
